# Peptide Barcodes for miRNA activity assessment in mammalian cells

**DOI:** 10.1101/2025.05.22.655559

**Authors:** Vasileios Cheras, Eirini Rousounelou, Jan-Lukas Aschenbach, Sven Panke, Yaakov Benenson

## Abstract

Studies of gene regulation require measurements of mRNA and protein levels of a regulated gene. Recently, parallel reporter assays have been introduced to study the regulation of multiple genes at once. While transcriptional regulation can be probed by next generation sequencing, post-transcriptional regulation requires the ability to measure multiple proteins in the same experiment. Multiplexing with the help of fluorescent proteins limits the addressable diversity of simultaneous measurements due to spectral overlap. Inspired by the utility of proteotypic peptides in targeted proteomics, here we show that genetically encoded peptide reporters (peptide barcodes) can be used to analyze multiple post-transcriptional pathways in parallel. We use RNA interference as an exemplary regulatory mechanism that occurs on both transcriptional and post-transcriptional levels. We measure the activity of multiple microRNAs in parallel using a peptide barcode-based miRNA sensor library. Fluorescent reporters are used to validate the accuracy of the miRNA activities reported via the peptide barcodes. Several assay optimization steps are explored leading to the robust activity profiling of nine miRNAs across three different cell lines. Overall, this study underlines the multiplexing potential of peptide barcodes to rapidly and quantitatively measure the post-transcriptional regulation.

## Introduction

Mammalian gene regulation is responsible for generating the phenotypic variation amongst cells sharing the same genotype, giving rise to a multicellular organism [1]. Elucidating the intricate regulatory pathways acting on the genomic, transcriptional and post-transcriptional level is key for understanding cellular differentiation and homeostasis [2] as well as disease phenotypes [3].

Over the past decade, massively parallel reporter assays (MPRAs) have been employed to establish the identity and dissect the function of putative *cis*-regulatory elements involved in gene expression, such as enhancers [4], promoters [5, 6], introns [7], and 3’ and 5’ untranslated regions (UTRs) [8, 9]. In these assays, each regulatory element variant exerts its effect on a reporter gene barcoded at the nucleotide level. Next generation sequencing allows for the identification and quantification of all the barcoded reporters in a single measurement, thus dissecting the functionality of every regulatory variant simultaneously [10]. The increased throughput of these methods has enabled functional characterization of millions of regulatory elements involved in transcriptional and post-transcriptional regulation [11].

Despite their great potential, MPRAs are limited to regulatory events that fully manifest themselves at the mRNA level. However, multiple regulatory mechanisms in cells are only fully observable at the protein level. RNA-binding proteins, miRNAs and ubiquitination-mediated protein degradation are a few examples of regulatory mechanisms that control protein levels independently of the abundance of the corresponding transcript [12–14]. Traditional methods to track multiple protein levels simultaneously are inherently low throughput because they rely, in some way, on optical readouts. These assays typically use conjugated antibodies, fluorescent probes or fluorescent protein fusions which suffer from spectral overlap and this limit the multiplexing capacity [15].

In contrast, mass spectrometry is much more amenable to multiplexing, as multiple proteotypic peptides can be detected simultaneously. These are unique peptide sequences representative of their proteins of origin, used to reliably identify and quantify them. Consequently, mass spectrometry enables the parallel analysis of hundreds of proteins in a single sample [16–18]. Similar to nucleic sequence barcodes, this capability inspired scientists to engineer peptide barcodes initially fused to the protein of interest but cleaved off prior to analysis [19, 20]. MS analysis of short barcodes is inherently simpler than the analysis of a full protein sequence, and in this fashion the barcodes serve as proxies for the protein to which they have been fused. In the past years, the peptide barcoding technology has enabled the identification of functional variants within large protein binder libraries within a single analysis step [20, 21], while it has also enabled the quantitative monitoring of up to 40 barcoded endogenous proteins involved in different pathways of *Saccharomyces cerevisiae* metabolism [22]. These applications promote peptide barcodes as a novel reporter class with increased diversity and orthogonality to conventional reporter proteins.

Based on these recent advances, in this study we asked whether the multiplexing potential of peptide barcodes can be exploited to monitor the multi-layered post-transcriptional regulation of the mammalian proteome. In particular, we focused on microRNA (miRNA) interference, a mechanism responsible for fine-tuning the levels of thousands of proteins in mammalian cells [23]. MiRNA regulation takes place on the mRNA level, where specific nucleotide sequences are recognized by the miRNA-RISC complex via base complementarity. The mRNA-bound RISC complex can promote mRNA degradation and/or translation inhibition manifesting its regulatory effect at the protein level [24]. As hundreds of miRNAs exist in mammalian cells, uniquely peptide barcoded reporter proteins can be potentially used to report on the regulatory effect of distinct miRNAs simultaneously.

As a proof of concept, we constructed a peptide barcode-based dual reporter system that can accurately quantify miRNA activity. We validated the functionality of the peptide barcode reporters by comparing the miRNA activity measured both via MALDI-TOF mass spectrometry (MALDI-TOF MS) and flow cytometry using traditional fluorescent reporter proteins. After identifying 18 functional and orthogonal peptides, we use them to create a miRNA sensor library that can quantify the activity of nine different miRNAs simultaneously. Further fine-tuning of the sensor design and the experimental procedure of the assay is explored leading to the simultaneous activity profile acquisition of nine miRNAs in three different cell lines. Overall, our results demonstrate that peptide barcodes can be used to study post-transcriptional regulation in a multiplexed fashion. Given the vast peptide barcode diversity, different regulatory mechanisms could therefore be monitored with increased throughput and multiplexing capabilities.

## Results

### Peptide-barcoded reporter assay design for miRNA activity assessment

Typically, fluorescent reporter-based miRNA sensors are used to assess the miRNA effect by measuring the reduction in the fluorescent reporter protein level resulting from miRNA-mediated inhibition [25–27]. Spectral overlap of fluorescent reporters limits the throughput of this approach to a one or two miRNAs per sample. To overcome this limitation, we surmised that cleavable peptide barcodes could be fused to conventional reporter proteins and, with the help of mass-spectrometry, dramatically increase throughput of assessing the activity of multiple miRNAs at once.

To this end we fused a unique peptide barcode sequence to the C-terminus of a fluorescent reporter protein regulated by a miRNA binding site in its 3’ untranslated region (UTR). To enable internal normalization of the readout, we also included a second, unregulated, reporter fused to a second (reference) peptide. The complete miRNA sensor is comprised of two expression cassettes driven by a constitutively active bidirectional cytomegalovirus (biCMV) promoter [Fig.1, A]. The promoter in the sense strand controls the transcription of an mCitrine activity reporter gene fused to a peptide barcode (activity peptide barcode), sensitive to miRNA-mediated regulation via four miRNA target sites in its 3’ UTR. The promoter in the antisense strand is responsible for the constitutive expression of a mCherry normalization reporter gene fused to another peptide barcode (normalization peptide barcode). The latter is used to account for both extrinsic and intrinsic factors that might bias the output of the bidirectional expression cassette, such as variations in the transfection efficiency and fluctuations in the expression machinery of the cell. The same 5’ UTR sequences are used in both sense and anti-sense directions to prevent any expression biases introduced by different 5’ UTR sequences. Additionally, a Kozak sequence is included in each expression cassette to ensure high reporter gene expression levels [28]. An N-terminal 6x-His-tag in each fusion reporter enables the separation from endogenous proteins using Ni-NTA-based affinity chromatography. Finally, a protease site encoded in a FLAG-tag encoded upstream of the peptide barcode allows for its proteolytic cleavage from the reporter protein prior to MS analysis using a site-specific protease.

**Fig. 1.**
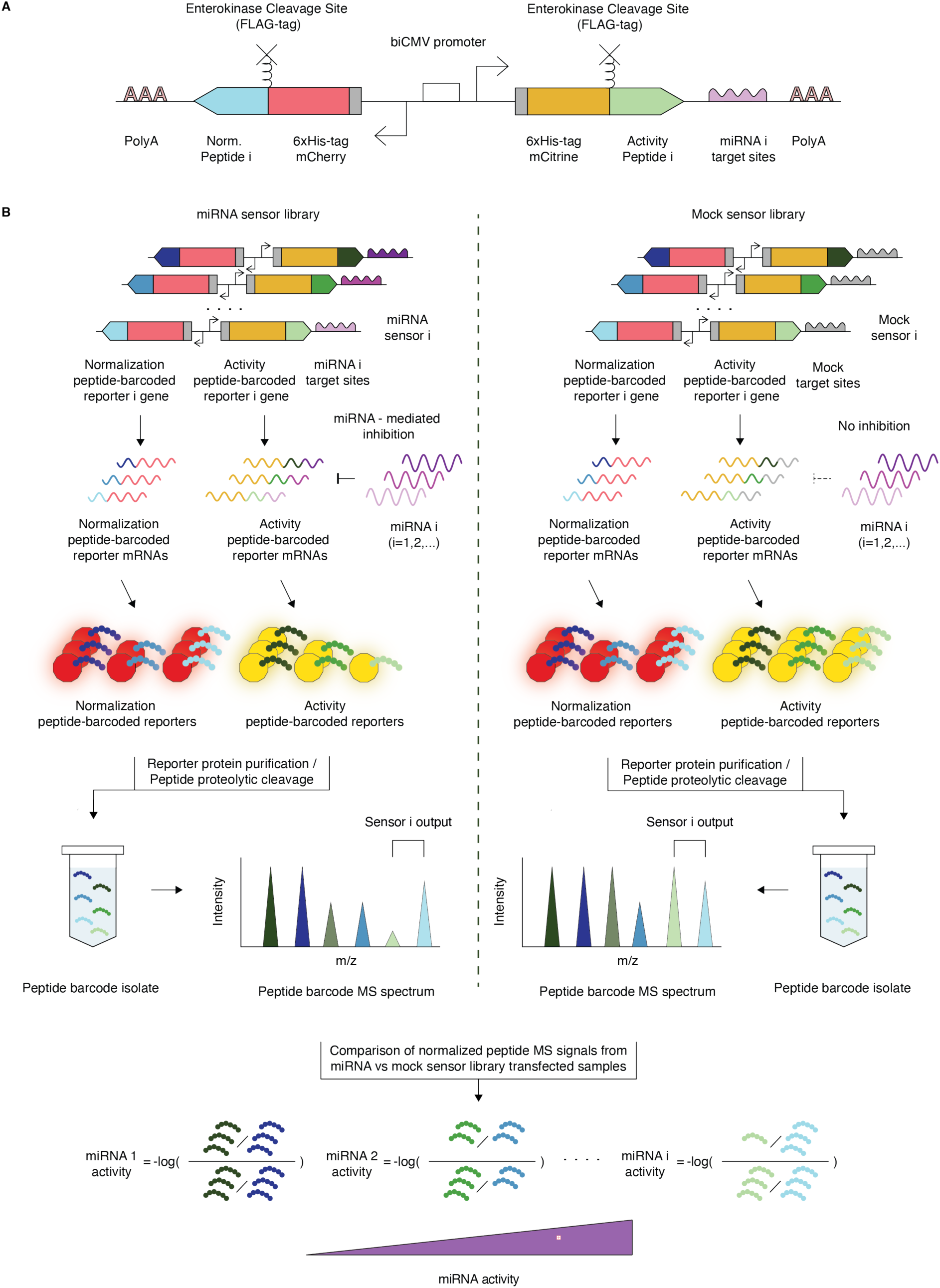
miRNA activity screening assay using peptide-barcoded miRNA sensors. **(A)** Genetic design of a peptide-barcoded miRNA sensor. The white box with opposing arrows depicts the biCMV promoter, driving the expression of the peptide-barcoded mCitrine and mCherry reporter genes. The grey boxes indicate the position of the sequence encoding the N-terminal 6x-His-tag. The peptide barcodes are attached to the C-terminus of each reporter protein and flanked by an enterokinase proteolytic cleavage site. The miRNA target sites encoded on the 3’ UTR are shown in purple. A polyadenylation signal is also located in the 3’ UTR of each fusion reporter gene. The inhibitory effect of endogenous miRNA i is exerted on the transcript encoding the peptide-barcoded mCitrine and carrying miRNA i complementary sites. **(B)** Workflow of multiplexed miRNA activity assessment using peptide barcodes as a readout. A library of miRNA sensor plasmids (left) or mock sensor plasmids (right) carrying sequences complementary to endogenous miRNAs (purple target sites) or to siRNA FF4 (grey-mock target sites), respectively, are transfected into cells, allowing for the expression of their peptide-barcoded activity and normalization reporter proteins under the presence (left) or absence (right) of miRNA inhibition. The cells are harvested and lysed, and the reporter proteins are affinity-captured on the surface of Ni-NTA resin via their N-terminal 6x-His-tag. After several washing steps, the activity and normalization peptides on the C-terminus of the captured reporter proteins are cleaved via enterokinase treatment. The cleaved peptide mix is separated from the immobilized reporter proteins, concentrated via lyophilization, and finally analyzed via mass spectrometry. For each sensor the relative ion abundance of its activity over normalization peptide is measured within the acquired spectrum. The ratio of the normalized activity peptides calculated in the presence (left) and absence (right) of miRNA inhibition yields the activity of a miRNA i.

A sensor library is created by assigning two unique peptide barcodes per miRNA target sequence. To calculate the activity of different miRNAs, two sensor libraries are tested separately: (i) a miRNA sensor library, where each sensor harbors the target site complementary to one miRNA of interest, which, if present, downregulates the abundance of the corresponding barcoded activity reporter via translational inhibition and/or mRNA degradation [29, 30] [Fig. 1, B, left]; (ii) a mock sensor library, where each sensor harbors a synthetic miRNA target site (FF4, [31]) lacking complementarity to any endogenous miRNA, thus allowing the non-repressed expression of barcoded activity reporters [Fig. 1, B, right]. Forty-eight hours after library transfection, cells are harvested and lysed, and all barcoded reporters are immobilized on a Ni-NTA resin. Subsequent on-resin proteolytic cleavage via a site-specific protease allows the isolation of peptide barcodes. The ion abundances of the activity and normalization peptide barcodes are acquired via MALDI-TOF, and a normalized reporter level is calculated for each sensor. Comparing the normalized peptide barcode levels stemming from cells transfected with the miRNA and mock sensor libraries, allows the deconvolution of each miRNA activity [Fig. 1, B bottom].

### Principles of peptide barcode design for quantitative MALDI-TOF measurements

Adequate peptide detectability and quantitation via MALDI-TOF MS are crucial for the applicability of the peptide barcode technology. To ensure these properties, specific rules were followed during the design of peptide amino acid sequences. Firstly, the designed peptide barcodes consist of 10 - 32 amino acids, spanning a molecular weight of 900 to 3,200 Da, thus benefiting from the good detectability of single-charged monoisotopic ions offered by MALDI-TOF MS in low m/z range [32]. Secondly, the m/z values of the peptides were separated from each other by a distance greater than 50 m/z units to avoid any sodium adducts skewing the signal of the peptides of interest. Thirdly, the designed peptide sequences lack any histidine, cysteine, or methionine residues to avoid oxidation and cross-linking events [20, 33]. Finally, a positively charged arginine or lysine residue was placed at the end of the amino acid sequence of each peptide for increased ionization efficiency [20, 33, 34]. An in-house algorithm generated a list of random peptide sequences fulfilling these criteria. A prediction algorithm was then used to evaluate the detectability of designed peptides *in silico*, and only peptides with detectability scores above the threshold were selected for further characterization [35].

Apart from the peptide sequence, sample preparation and acquisition have been shown to play a critical role in detectability. Heterogeneous sample crystallization and inadequate material ablation from the MALDI spot (sample coverage) have been previously linked to poor reproducibility of MALDI-TOF measurements [36]. At the same time, multiple shot accumulation has been shown to increase the detectability of low-abundance analytes [37]. Thus, ionizing material from multiple positions across a spotted sample and increasing the laser shot accumulation during spectrum acquisition was hypothesized to decrease measurement variability across technical replicates and improve peptide detectability. To test this hypothesis, we mixed equimolar amounts of two synthetic peptide barcodes (PepBC1 and PepBC2) and generated five technical replicate spots on a MALDI plate. These spots were ablated following a random pattern using an increasing number of accumulated laser shots and positions probed per spectrum, thus generating spectra of increased sample coverage [Fig. 2, A and Fig. S1 A-C]. Spectra with low number of accumulated laser shots resulted in highly variable absolute ion abundancies for both peptides (CV>40%). The measurement variation was merely improved when the relative abundance of the two peptides to each other was calculated (CV∼20%). However, a 40-fold increase in the accumulated shots acquired per spectrum dropped the technical replicate variation below 30% for absolute and below 6% for relative peptide abundance measurements.

**Fig. 2.**
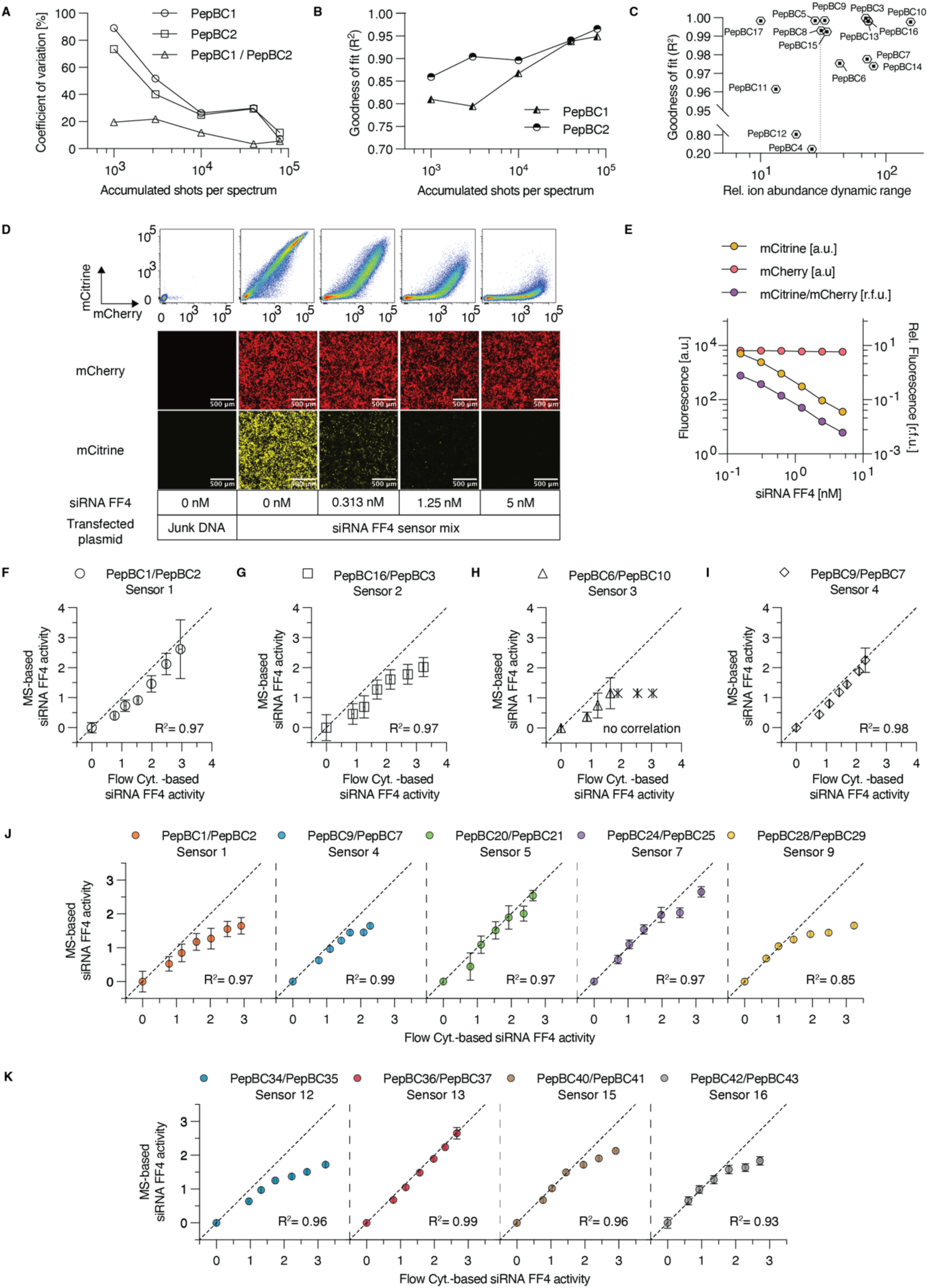
Validating peptide barcode detectability, functionality and multiplexing capabilities. **(A)** Examining the influence of increased sample coverage on the measurement of absolute and relative ion abundances of peptides in MALDI-TOF spectra. A line graph is depicted showing the effect of increasing accumulated shots (x-axis) on the coefficient of variation (y-axis) for absolute and relative ion abundance across technical replicate measurements. **(B)** Examining the influence of increased sample coverage on the standard curve construction for two synthetic peptide barcodes. Log-log lines were fitted to the standard curves of PepBC1 and PepBC2 acquired under different sample coverage settings. A line graph is depicted showing the effect of accumulated shots (x-axis) on the goodness of fit via the R^2^ (y-axis) for two standard curves of PepBC1 and PepBC2. **(C)** A scatter plot depicting the goodness of fit, and the dynamic range acquired from each peptide standard curve was constructed to evaluate the quantitative behavior of different peptides. A vertical dashed line was included to separate peptides exhibiting more than 30-fold signal reduction in their standard curves. **(D)** Flow cytometry plots (Top) and fluorescence micrographs (Bottom) illustrating the signals of peptide-barcoded mCitrine and mCherry reporters stemming from cells transfected with the four-member FF4-responsive sensor library and different concentrations of siRNA FF4 (0-5 nM). In the flow cytometry plots the mCitrine signal (D, y-axis) and the mCherry signal (D, x-axis) of single cells are depicted. The first column corresponds to cells transfected with junk DNA, lacking any fluorescent reporter expression. In the micrographs, mCitrine and mCherry-based reporters are represented by yellow and red pseudocolors, respectively. All micrographs are at 10x magnification. mCitrine: 300 ms exposure, LUT range of 0-65,535; mCherry: 300 ms exposure, LUT range of 0 – 65,535. Scale bars, 500µm. **(E)** Scatter plot depicting the average fluorescence values of mCitrine (yellow) and mCherry-based (red) reporters, as well as the average sensor output values (right y-axis, purple) across samples co-transfected with a four-member mock sensor library and increasing siRNA FF4 concentrations (0-5 nM) (x-axis). **(F-I)** Scatter plots comparing the siRNA FF4 activities acquired using flow cytometry measurements of individually titrated sensors (x-axis) or MALDI-TOF MS measurements of peptides isolated from the titrated sensor library (y-axis). Activity measurements, which were acquired individually via flow cytometry but could not be deconvoluted from the sensor library titration using MALDI-TOF MS, are depicted with an X mark. The dashed line corresponds to the line of identity in each graph. The R^2^ value corresponds to the result of the Pearson correlation test. **(J-K)** Scatter plots comparing the siRNA FF4 activities acquired using flow cytometry measurements of individually titrated sensors (x-axis) to the activities deconvoluted from MALDI-TOF MS measurements of peptide barcodes isolated from the titrated 15-sensor library (y-axis). The dashed line corresponds to the line of identity in each graph. The R^2^ value corresponds to the result of the Pearson correlation test.

Next, we tested how crucial is adequate sample coverage during sample acquisition to achieve quantitative MS measurements of peptide barcodes. For this, different synthetic peptide dilutions were measured under variable sample coverage conditions in the presence of a spike-in standard peptide. Indeed, increasing the number of accumulated shots per spectrum led to higher coefficient of determination (R^2^) values when a log-log line was fitted to the relative ion abundance measurements of the PepBC1 peptide titration curve [Fig. 2, B and Fig. S1, D]. Accumulation of 40,000 shots in 250 shot steps provided a robust titration curve fit (R^2^=0.94), while acquiring 80,000 shots per spectrum doubled the acquisition time per spot without improving dramatically the goodness of fit (R^2^=0.95). This observation was also confirmed for the titration curve of the designed peptide PepBC2 [Fig. 2, B and Fig S1, E]. After establishing the MALDI-TOF MS acquisition protocol, dilution series of fifteen additional designed peptides were individually tested, to identify more in-silico generated peptides with good quantitation potential. Thirteen peptides yielded standard curves with good fit (R^2^>0.95) in the concentration range of 0.021 μM – 1.7 μM, while eight of them had greater than 30-fold dynamic range of relative ion abundance [Fig. 2, C and Fig. S2, A-O].

The use of peptide barcodes to efficiently report varying levels of both the activity and normalization reporter proteins requires their analysis as a mixture. During the ionization of complex peptide mixtures, more efficient-to-ionize and more abundant peptides have been described to suppress the signal of other analytes, an effect known as ion suppression [38]. Thus, when measured with the optimized MALDI-TOF acquisition protocol, the selected peptide barcodes should retain their quantitation capacity even in the presence of more abundant peptides. To investigate this, we attempted to measure the PepBC1 and PepBC2 titration curves in a complex peptide background via MALDI-TOF MS. Specifically, we prepared mixtures of five peptides. In these samples, PepBC1 or PepBC2 concentration was varied while the rest of the peptides were present in constant concentrations. Both peptides retained a comparable goodness of fit and dynamic range of detection when tested in the complex mix. The ion abundance of peptide barcodes with a constant concentration in the mix remained steady throughout measurements [Fig. S1, F-G]. This result confirmed that ion suppression did not affect the detectability of PepBC1 and PepBC2 in the peptide mix. Thus, we speculated that other individually titrated peptides would also retain their dynamic range of detection when analyzed simultaneously. After establishing a MALDI-TOF MS acquisition protocol that allows for relative quantification of *in silico*-generated peptide barcodes with good quantitation potential, we moved on to the design of peptide-barcoded sensors for determining miRNA activity in mammalian cells.

### Peptide-barcoded sensor functionality and multiplexing capabilities validation

To prove the functionality and multiplexing capabilities of the peptide-barcoded reporters, a small sensor library was created by mixing four sensors. These were equipped with eight peptide barcodes, which exhibited good quantitation potential when titrated as synthetic peptides. Four repeats of an FF4 target sequence were used in each sensor’s sense 3’ UTR to render them responsive to externally provided siRNA FF4 but unresponsive to endogenous miRNAs [27, 39]. The library was co-transfected with increasing amounts of siRNA FF4 into HEK293 cells to titrate the response of all four sensors simultaneously. Flow cytometry and fluorescence microscopy analysis confirmed the decreasing cumulative fluorescent signal of mCitrine activity reporters with increasing concentrations of siRNA FF4, while the cumulative signal of mCherry-based normalization reporters remained unchanged [Fig.2, D-E].

MS analysis of the isolated peptide barcodes confirmed a decrease in all four sensor outputs along the siRNA FF4 concentration gradient [Fig. S3, A]. To evaluate the siRNA FF4 repression strength, the response of each sensor was normalized to its output in the non-repressed state (0 nM siRNA FF4). The normalized responses were log-transformed to calculate siRNA activities acquired via MS, which subsequently were compared to those derived from individually titrated sensors via flow cytometry [Fig.2, F-I]. The responses of sensor 1 (R^2^=0.97), sensor 2 (R^2^=0.97), and sensor 4 (R^2^=0.98) significantly correlated between the two measurement methods. The activity peptides of sensor 1 (PepBC1), sensor 2 (PepBC16) and sensor 4 (PepBC9) were detectable even under strong siRNA-mediated repression in the transfected sample (5 nM siRNA FF4). The higher activity standard deviations acquired by the peptides of sensor 2 were justified by the lower ion abundance (∼10^3^ a.u.) of the normalization peptide PepBC3 in comparison to the other normalization peptides (∼10^5^ a.u.) that cannot reliably normalize any abundance biases of the activity peptide PepBC16 [Fig. S3, A]. Interestingly, the fluorescence signal of the fusion mCherry-PepBC3 reporter was lower compared to that of other peptide-barcoded mCherry reporters, indicating low expression of this reporter protein [Fig. S3, B]. The MS output of sensor 3 could not be measured when 1.25 nM siRNA FF4 was used, due to low PepBC6 detectability, thus limiting the utility of the sensor [Fig. S3, A]. A closer examination of the spectra regions of PepBC3 and PepBC6 revealed the presence of several impurity peaks that potentially contributed to signal suppression for both peptides [Fig. S3, C-D]. The same impurity peaks were detected after analyzing peptide isolates from cells lacking any peptide barcode expression, suggesting that they arise from cell culture impurities carried over during protein purification [Fig. S3, E-F].

Sensor titration using siRNA FF4 represents a straightforward way to characterize the peptide barcode response, considering biases neglected during the construction of synthetic peptide barcode standard curves, such as differences in the fusion reporter expression and carry-over impurities during peptide purification. Thus, we hypothesized that we could utilize the sensor titration assay to functionally characterize multiple peptide-barcoded sensors simultaneously without previously testing the MS detectability of synthetic peptide barcodes. To expand the peptide-barcoded sensor repertoire, 24 additional peptides with good *in silico* detectability scores were designed to generate 12 new mock sensors. These sensors were combined with the previously chosen three sensors to create a 15-member mock sensor library, which was titrated using increasing siRNA FF4 amounts in HEK293 cells. The peptide ion abundances corresponding to each sensor were used to calculate the sensor’s response [Fig. S4, A-O], which was then converted to siRNA FF4 activity. In parallel, individual sensor titrations were performed to acquire the flow cytometry-based siRNA activities. Those values were then compared to the MS-based activity measurements acquired from the 15-sensor library transfection [Fig.2, J-K & Fig. S5]. Ten sensors responded to siRNA FF4 concentrations equal to or greater than 2.5 nM [Fig. S4, A-D, F, H, K, L, N, O], exhibiting strong peptide MS signals. The remaining five sensors were discarded, as their activity peptide barcodes were non-responsive to siRNA concentrations equal or less than 0.625 nM [Fig. S4, E, G, I, J, M]. As previously observed, the normalization peptide PepBC3 of sensor 2 exhibited low ion abundances and was thus excluded from further usage [Fig. S4, B]. The increasing siRNA FF4 activity reported by the nine functional mock sensors significantly correlated with the activity determined via flow cytometry of individually titrated sensors [Fig.2, J-K].

### miRNA activity profiling of three mammalian cell lines

After identifying nine functional peptide-barcoded sensors, we asked if they could be used to capture the activity profile of nine endogenous miRNAs across three different cell lines, namely HEK293, HeLa, and HuH-7. The choice of miRNAs was based on previous miRNA activity screens [25, 40, 41], as well as miRNA expression levels measured in-house. The nCounter miRNA profiling platform was used to indirectly estimate the relative miRNA abundance levels across the three different cell lines [Fig. S6] [42]. The final panel included miRNAs of diverse activities and abundances that create a profile characteristic of each cell line [Fig. 3, A]. The peptide-barcoded miRNA sensors were created by exchanging the FF4 targets with the complementary target sites of the chosen miRNAs. Each sensor was first transfected in isolation and compared to its siRNA FF4-responsive counterpart to verify the activity profile of all three cell lines via flow cytometry [Fig. 3, A (bottom)]. The sensors responded to the different endogenous miRNAs, detecting repression levels of up to 2.5 orders of magnitude, with miR-17, miR-21, and miR-122 showing the highest activities in HEK293, HeLa, and HuH-7 cells, respectively, in accordance with the literature [40, 43]. When comparing miRNA activity to the relative miRNA abundance measurements, only a weak correlation was found across all cell lines (R^2^=0.34). It is evident that miRNAs of the same expression level do not necessarily result in the same protein downregulation [Fig. 3, B]. This finding confirms the complex nature of miRNA regulation, highlighting the necessity for activity measurements at the protein level.

**Fig. 3:**
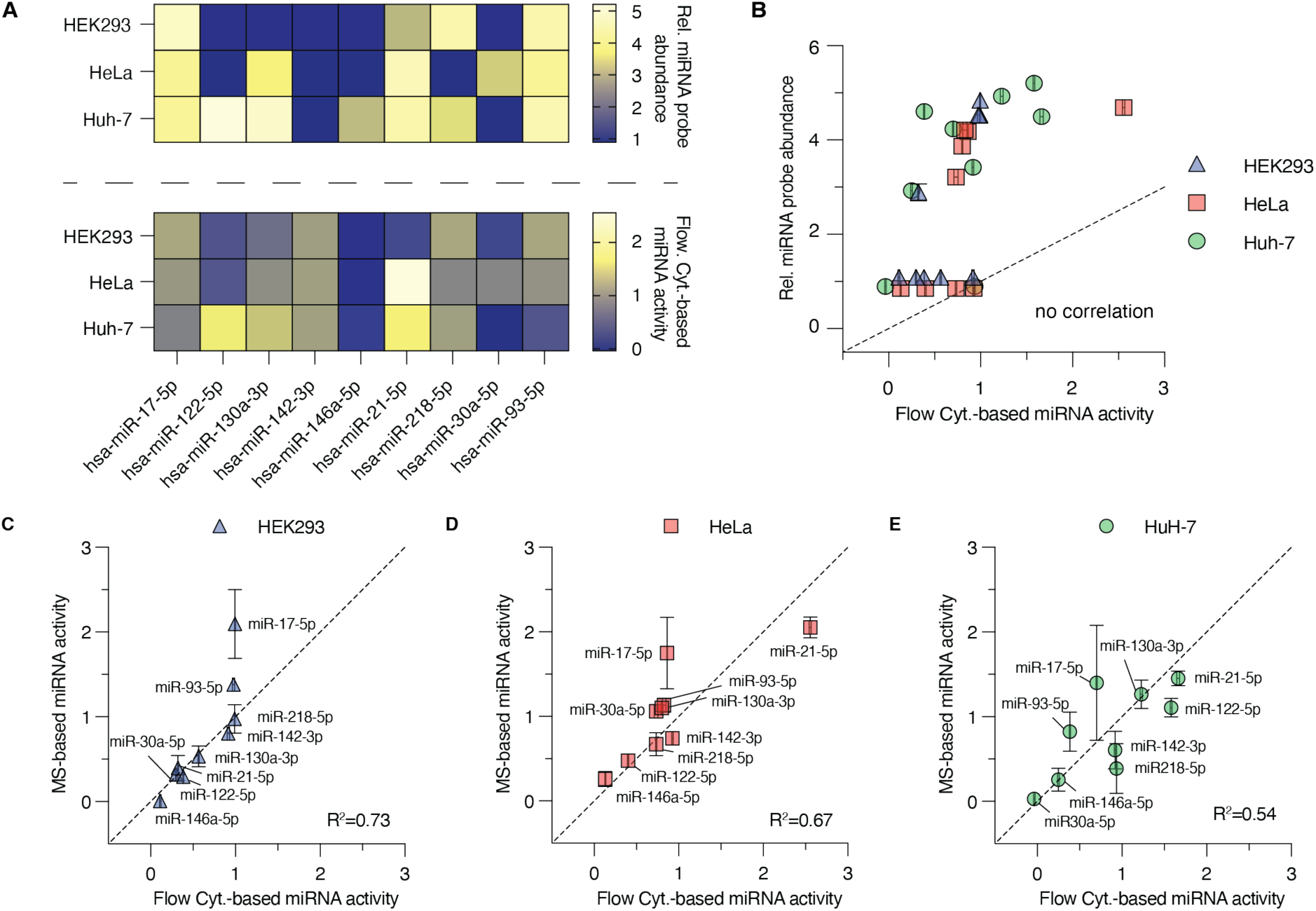
miRNA abundance and activity profiling of HEK293, HeLa and HuH-7 cells. **(A)** (Top) Heatmap depicting the relative abundance of miRNA probes detected in HEK293, HeLa, and HuH-7 cells, corresponding to the nine miRNAs selected for miRNA activity cell line profiling. (Bottom) Heatmap depicting the activity of selected miRNAs measured via flow cytometry of individually transfected miRNA sensors. **(B)** Scatter plot comparing the relative abundance of miRNA probes (y-axis) to the miRNA activity acquired via flow cytometry (x-axis) for HEK293 (blue), HeLa (red), and HuH-7 cells (green). The results of the Pearson correlation test between miRNA probe relative abundance and activity across all cell lines, as well as the dashed identity line are depicted on the graph. **(C-E)** Scatter plots comparing the activities of nine miRNAs in HEK293 (A), HeLa (B), and HuH-7 (C) cells, acquired using flow cytometry measurements of individually titrated sensors (x-axis) or MALDI-TOF MS measurements of peptides isolated from the titrated sensor library (y-axis). The dashed line corresponds to the line of identity in each graph. The R^2^ value corresponds to the result of the Pearson correlation test.

In the next step, the three cell lines were transfected with the mock sensor and miRNA sensor libraries separately to acquire their respective miRNA activity profiles via MS. The activity profiles were compared to those acquired via flow cytometry of individual sensor transfections [Fig. 3, C-E]. The correlation between the two methods was adequate for HEK293 (R^2^=0.73) and HeLa cells (R^2^=0.67) but weak for HuH-7 cells (R^2^=0.54). Moreover, the MS-derived measurements from HuH-7 cells exhibited large standard deviations, increasing the uncertainty of the profiling results [Fig. 3, C-E]. Examination of the peptide barcode MS signals of mock sensor libraries revealed similar peptide ion abundances for HEK293 and HeLa cells but yielded lower values in Huh-7 samples [Fig. S7, A]. The larger standard deviations observed with HeLa and HEK293 samples are connected to biases during spectra acquisition that were balanced by the signal of the normalization peptides, resulting in similar mock sensor output levels [Fig. S7, B]. On the contrary, low peptide detectability led to lower sensor outputs in HuH-7 samples [Fig. S7, B], and that subsequently reduced the accuracy of the acquired miRNA activities in this cell line [Fig. 3, E].

### Accuracy improvement of Ni-NTA based miRNA activity profiling

Given the promising results of the miRNA profiling campaign, our efforts focused on optimizing several experimental parameters to ensure robust peptide signals across all cell lines tested. Firstly, we observed differences in protein expression, as HuH-7 and HeLa cells yielded less reporter protein output than HEK293 cells, despite being transfected with equal plasmid amounts [Fig. S8, A]. Thus, transfection optimization was necessary to adjust the reporter expression levels across different cell lines [Fig. S8, B-C]. Secondly, the m/z ratio for half of the peptides falls in a part of the spectrum (m/z 600-2,000) in which many impurities from the purification process were present. Such impurities can originate from proteolytic products of endogenous proteins non-specifically bound to the Ni-NTA resin [Fig. S8, D-F] or Tween 20 surfactant from the lysis solution [Fig. S8, G]. To remove these biases, new peptide barcodes were designed with tailored molecular weights to avoid noisy parts of the spectra. These were combined with peptide barcodes previously exhibiting strong MS signals across the different cell lines, thus generating an optimized peptide barcode repertoire. In addition, a low molecular weight surfactant replaced Tween 20 to minimize any surfactant-related impurity peaks influencing peptide detectability [Fig. S8, H]. Finally, halving the spotted sample amount on the MALDI plate accelerated the sample evaporation, rescuing the sub-optimal sample crystallization that could contribute to the large standard deviations across technical replicate MS measurements [44].

Considering all these improvements, a nine-member sensor library was created by combining eight newly designed peptide-barcoded miRNA sensors (sensors 17-24) with sensor 13 (PepBC36/PepBC37), which was retained due to its robust output across cell lines [Fig. S7, A-B]. This optimized set of sensors was equipped with FF4 targets and titrated using siRNA FF4 in HuH-7 cells, as this cell line had proven the most difficult to profile. All the isolated peptides were detectable across the siRNA FF4 concentration range (ion abundances >10^4^ a.u.), while the sensor outputs decreased with increasing siRNA FF4 levels [Fig. S9, A-I]. The response curves of sensors 23 and 24 were less steep compared with those of other sensors [Fig. S9, H, I]. Nonetheless, for all nine sensors the siRNA FF4 activity acquired via MS correlated significantly with the flow cytometry-derived measurements of individually titrated mock sensors (R^2^>0.84) [Fig. 4, A-B].

**Fig. 4:**
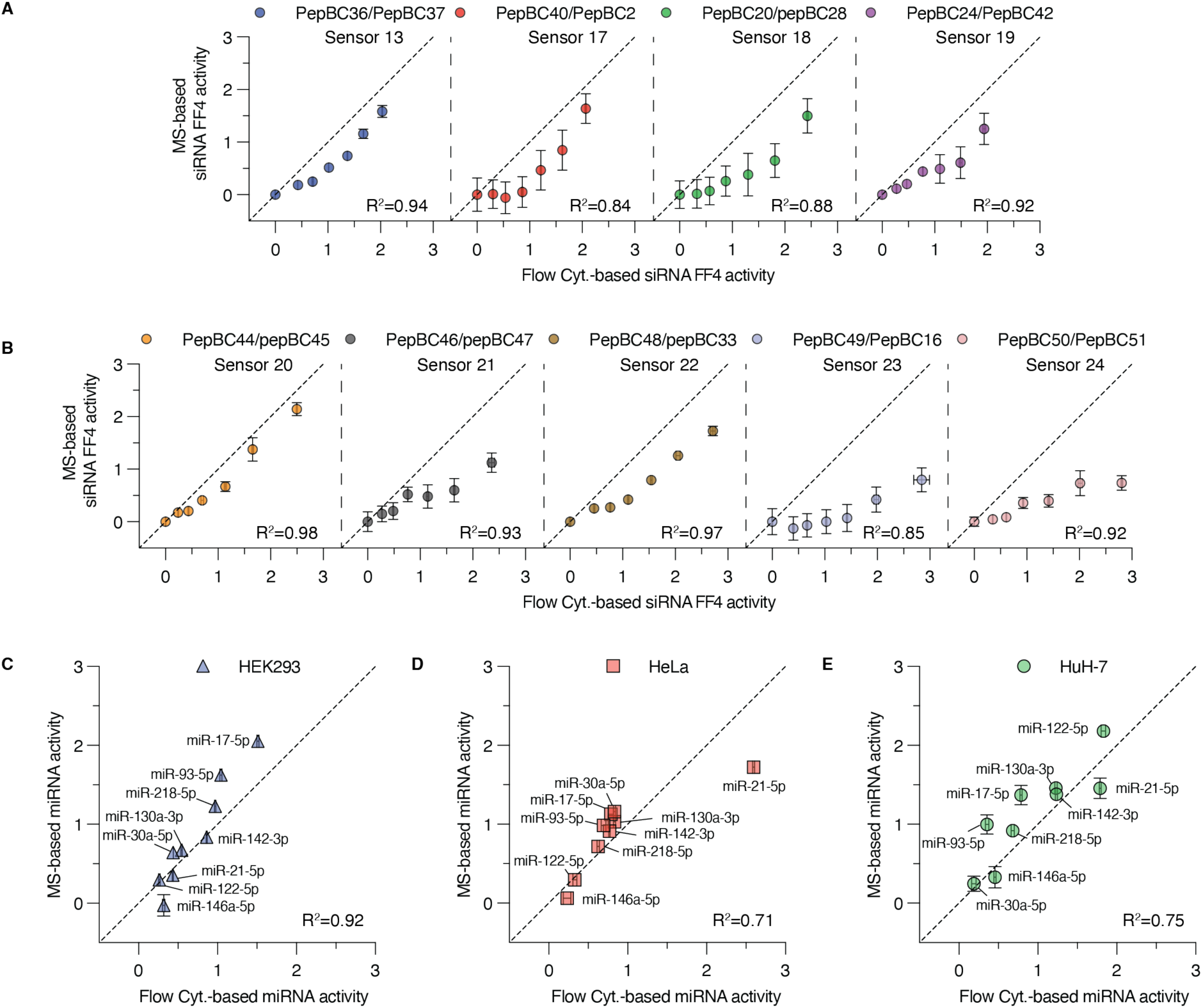
Optimized Ni-NTA-based miRNA activity assessment. **(A-B)** Scatter plots comparing the siRNA FF4 activities acquired using flow cytometry measurements of individually titrated sensors (x-axis) or MALDI-TOF MS measurements of peptides isolated from the titrated sensor library (y-axis). **(C-E)** Scatter plots comparing the activities of nine miRNAs in HEK293 (C), HeLa (D), and HuH-7 (E) cells, acquired using flow cytometry measurements of individually titrated sensors (x-axis) or MALDI-TOF MS measurements of peptides isolated from the titrated sensor library (y-axis). The dashed line corresponds to the line of identity in each graph. The R^2^ value corresponds to the result of the Pearson correlation test.

Next, the miRNA activity profiling of the three cell lines was repeated with the optimized set of miRNA sensors. In contrast to the previous profiling attempt [Fig. 3, C-E], the correlation between MS-acquired and flow cytometry-acquired miRNA activity profiles was significant across all three cell lines (HEK293 (R^2^=0.92), HeLa (R^2^=0.71), HuH-7 (R^2^=0.75)). At the same time, the standard deviation of MS measurements was drastically reduced [Fig. 4, C-E]. All peptides of the mock sensor library yielded robust signals across the different cell lines [Fig. S9, J]. Peptide isolates from HuH-7 cells exhibited higher ion abundance levels than peptides expressed in HeLa cells; however, peptides isolated from HEK293 cells retained the highest MS signals. Overall, the new experimental conditions combined with the optimized sensor set led to significantly higher peptide signals compared to the initial profiling attempt [Fig. S9, K].

### Impurity decrease further improves miRNA activity profiling

Despite all optimization efforts, multiple unidentified peaks were still present in the analyzed spectra [Fig. S8, D-F]. These impurities compete with the peptides of interest for ionization energy during sample ablation, thus hindering the detection of low-abundance peptides, such as activity peptides under strong miRNA regulation. Ni-NTA-based protein purification is reportedly prone to carry over of endogenous proteins and serum proteins from the culture medium, exhibiting affinity towards nickel ions [45]. Thus, we speculated that using a more selective purification method should yield protein isolates of higher purity. The presence of a FLAG-tag in the fusion reporter proteins, serving as a proteolytic cleavage site, allowed us to use an anti-FLAG resin to selectively immunoprecipitate the peptide-barcoded reporters of miRNA or mock sensors. Lysates from HEK293, HeLa, and HuH-7 cells lacking any peptide-barcoded reporter expression underwent anti-FLAG-based affinity purification followed by enterokinase treatment to evaluate the background spectra purity of the different cell lines [Fig S10, A]. The total number of peaks detected was significantly lower in anti-FLAG-purified than in Ni-NTA-purified peptide isolates for all tested cell lines [Fig S10, B]. The impurity peak reduction was evident across most spectrum m/z regions corresponding to peptide barcodes, as proven by the significantly reduced background summed intensities measured in these regions [Fig S10, C]. Removing impurities competing for ionization energy in the MS resulted in a significantly increased MS signal for the PepBC19 peptide that was spiked in a HEK293 peptide isolate [Fig S10, D].

This observation led to the hypothesis that reducing carry-over impurities could increase the MS signal of the peptide barcodes and consequently improve the miRNA activity measurements. Thus, the optimized miRNA and mock sensor libraries were transfected into the previously profiled cell lines, but this time followed the anti-FLAG purification protocol. Surprisingly, the peptide barcode ion abundances derived from HEK293 and HuH-7 cells transfected with miRNA or mock sensor libraries were lower than those acquired in the previous profiling attempt following the Ni-NTA purification strategy [Fig. 5, A-B]. HeLa-derived peptide isolates exhibited slightly stronger MS-signals when the anti-FLAG protocol was followed for purification. However, this observation was justified by the higher cell transfection efficiencies in the anti-FLAG-based-versus the Ni-NTA-based profiling campaign, thus biasing the isolated peptide barcode amounts [Fig. S11, A-B].

**Fig. 5:**
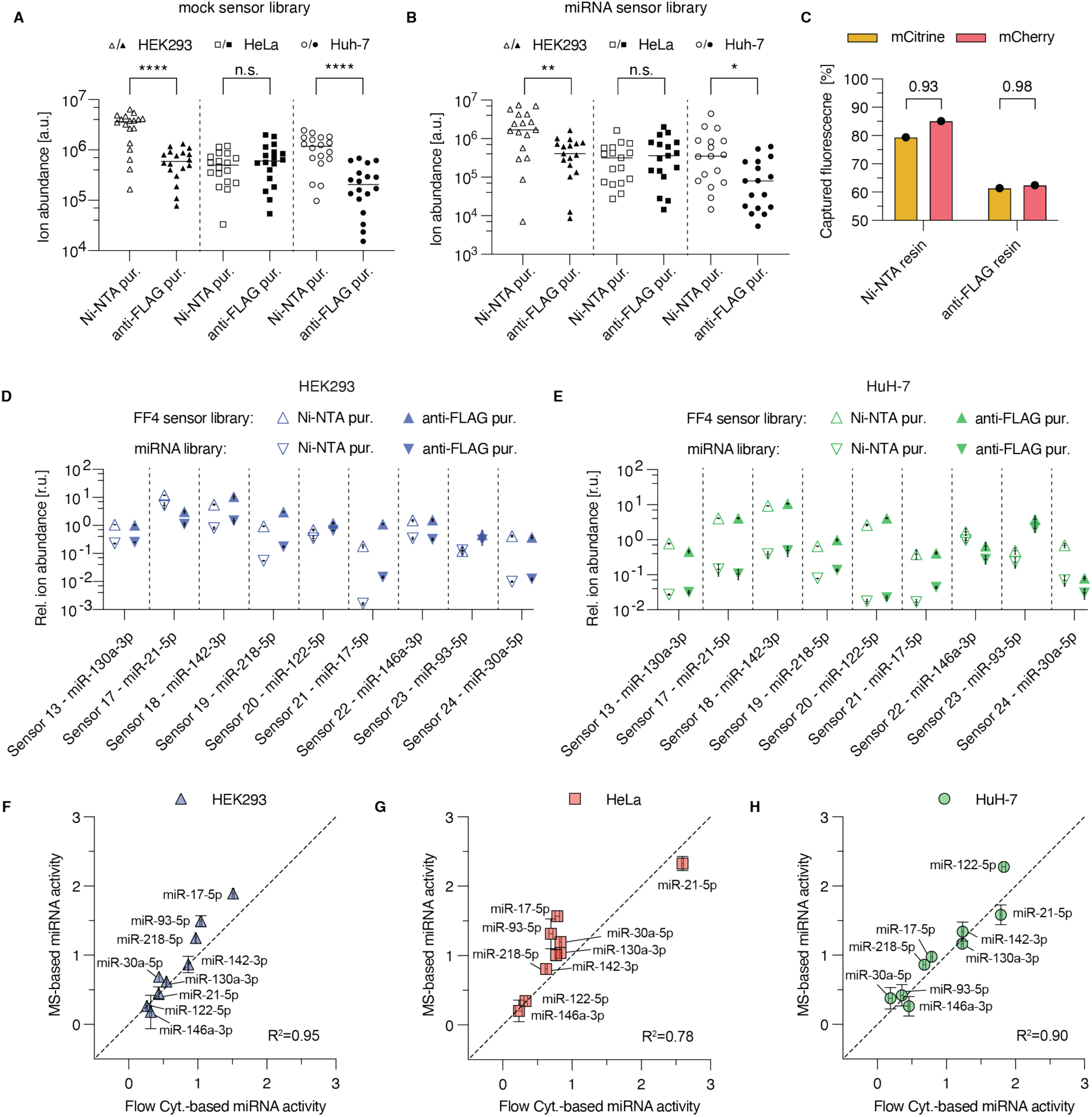
Anti-FLAG-based miRNA activity assessment. **(A-B, D-E)** Comparison of peptide barcode ion abundances (A-B) and sensor outputs (D-E) acquired from HEK293 (blue) or HuH-7 (green) cells transfected with a mock miRNA library (triangles pointing down) or a miRNA sensor library (triangles pointing up) subjected to either Ni-NTA (no fill symbols) or anti-FLAG (filled symbols) purification. Statistical significance of the differences observed across Ni-NTA-vs anti-FLAG-purified samples was calculated using an unpaired two-tailed Welch’s t test (A-B) (*p < 0.05; **p < 0.01; ****p < 0.0001). The dashed lines in A and B are separating the comparison across HEK293 (triangle) and HuH-7 cells (circle). The dashed lines in D and E are separating the outputs of different sensors. **(C)** Bar plot depicting the percentage of mCitrine (yellow) or mCherry (red) fluorescence captured on Ni-NTA and anti-FLAG resins from HEK293 cell lysates transfected with a mock sensor library. The mCitrine/mCherry fluorescence ratios are depicted on top of the bars. For this experiment, the initial set of mock sensors was used. **(F-H)** Scatter plots comparing the activities of nine miRNAs in HEK293 (F), HeLa (G), and HuH-7 (H) cells, acquired using flow cytometry measurements of individually titrated sensors (x-axis) or MALDI-TOF MS measurements of peptides isolated from the titrated sensor library (y-axis). The dashed line corresponds to the line of identity in each graph. The R^2^ value corresponds to the result of the Pearson correlation test.

These results indicated that lower peptide barcode amounts were isolated following the anti-FLAG purification protocol. Indeed, analyzing the fluorescent reporter signal captured by either Ni-NTA or anti-FLAG resins from lysates of HEK293 cells transfected with mock sensors revealed that the Ni-NTA resin captured 79.4% of the mCitrine- and 85.2% of the mCherry-based reporters in the lysate, while the anti-FLAG resin retained only 61.4% and 62.4%, respectively [Fig. 5, C]. Despite the lower binding capacity of the anti-FLAG resin, the high purity of peptide isolates still resulted in sensor outputs comparable to the measurements acquired from Ni-NTA-purified samples [Fig. 5, D-E]. Interestingly, the ratio between mCitrine- and mCherry-based peptide fusion reporters captured was slightly higher for the anti-FLAG resin (0.98), indicating that a higher percentage of mCitrine fusion reporters can be purified compared to the Ni-NTA resin (0.93) [Fig. 5, C]. Higher levels of mCitrine-based reporters correspond to increased representation of the miRNA-regulated peptide barcodes in the peptide isolate, contributing to stronger sensor outputs in the MS. This observation serves as an explanation for the higher sensor outputs in anti-FLAG purified samples compared to Ni-NTA purified samples [Fig. 5, D-E]. Finally, the MS-based miRNA activity measurements of anti-FLAG-purified peptide isolates strongly correlated with those acquired from individual sensors tested via flow cytometry [Fig. 5, F-H]. In fact, the correlation was improved in comparison to the previous Ni-NTA-based profiling for all three cell lines (HEK293 (R^2^=0.95), HeLa (R^2^=0.78), HuH-7 (R^2^=0.90)), thus showing that higher purity peptide barcode mixes can provide good quantitative results with smaller peptide quantities.

## Discussion

In the past decades, several gene reporter assays have been developed to study gene expression regulation [46, 47]. Next-generation sequencing technology enabled these assays to rapidly and simultaneously characterize a plethora of genetic elements regulating the transcriptome [9, 48–51]. However, studying gene expression at the protein level is characterized by lower throughput, due to the reduced orthogonality of the optical readout reporters used in conventional fluorescent or luminescent reporter assays [52, 53]. So far, the Sensor-seq platform proposed by Mullokandov *et al* is the only gene reporter assay that allowed the parallel categorization of hundreds of miRNAs into activity bins using only two reporter proteins and DNA barcodes [41]. Nevertheless, the increased assay throughput comes at the expense of precise determination of each miRNA activity. In this study, we combined miRNA activity profiling via the peptide barcoding technology to MALDI-TOF MS analysis to create a novel gene reporter assay that can simultaneously assess the activity of nine different miRNAs across three different cell lines. To the best of our knowledge, the proposed assay is the first to accurately, quantitatively and simultaneously assess the activity of multiple miRNAs at single miRNA sensor resolution.

The proposed miRNA activity assay relies on monitoring the signal decrease of the peptide barcodes between samples transfected with miRNA and mock sensor libraries. Thus, maximizing the peptide signal in the mass spectrometer was crucial to provide robust sensitivity across different miRNA-mediated repression levels. Throughout the assay development, several factors were identified to affect the MS detectability of the peptide barcodes. Firstly, the amino-acid selection on each peptide sequence resulted in distinct ionization efficiencies in accordance with the literature. Additionally, the peptide barcoded reporter proteins showed different fluorescence levels, suggesting that each peptide barcode influenced also the fusion reporter expression. Furthermore, the peptide barcode MS signals was proven to be rather sensitive to suppression ionization effects from abundant carry-over impurities across the different purification steps. All these factors translated into variable responses for each miRNA sensor, limiting the robustness of the proposed assay. Several steps, such as adapting the peptide barcode design, optimizing the reporter protein expression, switching to a more selective purification method, as well as improving the MS sample preparation and acquisition mitigated the limitations experienced during assay development.

Taking advantage of the fast miRNA activity profiling, the genetic design of the peptide-barcoded miRNA sensors could be adapted to elucidate determinants of miRNA-mediated regulation, such as: (i) the position of miRNA target sites in the mRNA transcript; (ii) the miRNA target site architecture and complementarity; or (iii) combinatorial regulation by different miRNAs. Furthermore, peptide-barcoded sensors could also be employed to study multiplexed regulation, as different cis-regulatory elements, miRNA target sites and even degradation tags can be combined in the sensor design and peptide barcodes can then be used to evaluate the combinatorial effect of both transcriptional and post-transcriptional regulation at the protein level.

Differential regulatory element profiles are of particular interest, as they characterize different normal and diseased cell types. These profiling data hold great value for developing sophisticated synthetic gene circuit therapeutics, which depend on accurate regulatory element characterization to specifically act on targeted cell populations [40, 54]. Thus, the presented assay could potentially be exploited to profile any cell line in vitro rapidly and accurately requiring only a limited number of experimental samples. As computational tools have automated the design of such cell classifiers [55, 56], one can envision that large profiling campaigns using peptide-barcoded sensor libraries could rapidly generate data, which in turn are used to design cell classifier circuits for many different indications. As in vivo and in vitro profiling data have been reported to differ significantly, it is important to be able to profile biologically relevant samples [25]. With minor adaptations, the proposed assay can utilize virus-based libraries of peptide-barcoded sensors that can transduce and profile clinically relevant samples, such as primary cells or xenografted tumors in mice. The expanded applicability of the assay is further facilitated by its increased throughput, which reduces the biological material needed for profiling compared to conventional fluorescence-based reporter assays.

Overall, further expansion of the peptide barcode repertoire and optimization of the experimental protocols, shall boost the multiplexing capacity of the proposed reporter assay. In the future, peptide barcode-based assays in parallel with NGS-based MPRAs hold the potential to elucidate the complex landscape of gene expression regulation, providing invaluable insights for both applied and basic research.

## Supporting information

Table_S10_Acquisition_parameters_used_for_MALDI_TOF_measurements

Table_S11_S12_S13_S14_Analysis_parameters_used_for_MALDI_TOF_and Flow_Cytometry_measurements

Table_S15_Statistical_analysis_information_per_figure

Tables S1_S2_Peptide_barcode_properties_and_peptide_barcoded_sensors

Tables S3_S4_S5_S8_S9_primers_DNA_fragments_siRNAs

Tables S6_S7_Plasmid_Cloning_procedure_and_plasmid_sequences

## Acknowledgements

The project was funded by the NCCR “Molecular Systems Engineering”, SNF Grant 175760, and ETH Zurich. We would like to thank the Single Cell Unit of D-BSSE for help with flow cytometry and microscopy; Dr. Todd Duncombe and Simon Berlanda from the lab of Prof. Dr. Petra Dittrich for helping with the MALDI-TOF measurements; Dr. Matteo Lampis for helping with the Nanostring miRNA profiling experiment as well as its MATLAB analysis; Dr. Tania Roberts for manuscript curation; Dr. Bartolomeo Angelici, Dr. Jiten Doshi, Dr. David Schweingruber, Dr. Phillip Müller-Thümen, Dr. Gabriel Senn and other Benenson lab members for discussions.

## Author Contributions

V.C. conceived research, performed most of the experiments, analyzed data, and wrote the paper. E.R. generated the peptide barcode amino-acid sequences, performed the synthetic peptide quantification, and helped with the first miRNA activity profiling of the cell lines. J.L.A. established and tested the anti-FLAG based peptide purification and helped with the last miRNA activity profiling of the cell lines. S.P. and Y.B. conceived research, wrote the paper, and supervised the project.

## Methods

Key resources table

| REAGENT or RESOURCE | SOURCE | IDENTIFIER |
| --- | --- | --- |
| Bacterial and virus strains |  |  |
| *Escherichia coli* TOP10 | ThermoFisher | Cat#C404010 |
| Chemicals, peptides, and recombinant proteins |  |  |
| Sphero Rainbow Calibration Particles 8 peak beads | Spherotech | Cat#PCP-30-5A |
| Lipofectamine 2000 | ThermoFisher | Cat#11668019 |
| Triton X-100 | AppliChem | Cat# A1388.0500 |
| Tween 20 | Sigma-Aldrich | Cat# P9416-100ML |
| Ni-NTA agarose | QIAGEN | Cat# A1388.0500 |
| Imidazole | Sigma-Aldrich | Cat# 56749-50G |
| Enterokinase | NEB | Cat# P8070L |
| α-cyano-4-hydroxycinnamic acid powder | Sigma-Aldrich | Cat# 70990 |
| Gibson Assembly Master Mix | NEB | Cat# E2611S |
| Synthetic peptides used, see Table S1 | This paper | N/A |
| Critical commercial assays |  |  |
| ZR Plasmid Miniprep - Classic | Zymo Research | Cat# 4016 |
| PureYield™ Plasmid Midiprep System | Promega | Cat# A2495 |
| Norgen Endotoxin Removal Kit Midi | Norgen Biotek | Cat# 52200 |
| miRVana miRNA Isolation kit | ThermoFisher | Cat# AM1560 |
| DYKDDDDK Fab-Trap Agarose Kit | ChromoTek | Cat# ffak-20 |
| nCounter miRNA expression assay | Nanostring | N/A |
| Experimental models: Cell lines |  |  |
| HEK293 (293-H) | Invitrogen | Cat# 11631-017 |
| HeLa | ATCC | ATCC Cat# CCL-2;<br>RRID:CVCL_0030 |
| HuH-7 | JCRB | Cat# JCRB0403;<br>RRID:CVCL_0336 |
| Oligonucleotides |  |  |
| Human miRNA-tags | Nanostring | Cat# Hu-MIRTAG-12 |
| Human miRNA CodeSet | Nanostring | Cat# CSO-MIR3-12 |
| Primers, see Tables S3, S4 and S8 | This paper | N/A |
| Gene fragments/synthetic DNA sequences/gBlocks,<br>see Table S5 | This paper | N/A |
| miRIDIAN negative control #2, see Table S9 | Dharmacon | Cat#CN-002000-01-05 |
| siRNA-FF4, see Table S9 | Dharmacon | N/A |
| Recombinant DNA |  |  |
| pVC001 | Genewiz | N/A |
| pKH025(Ef1a-mCitrine) | Prochazka et al.,2014<br>[57] | N/A |
| pKH026(Ef1a-mCherry) | Prochazka et al.,2014<br>[61] | N/A |
| pBH265 (junkDNA) | Haefliger et al.,2016<br>[26] | N/A |
| pRA52 | Altamura et al.,2022<br>[58] | N/A |
| pMD26 | Dastor et al., 2018 [25] | N/A |
| pEF1/His B | Invitrogen | Cat# V92220 |
| pSelect His-Tag | InvivoGen | Cat# psetb-chis |
| pEZYmyc-His | Guo et al., 2008[59] | Addgene Plasmid 18701 |
| Additional plasmids, see Tables S2, S6 & S7. | This paper | N/A |
| Software and algorithms |  |  |
| In-house MS analysis Matlab based algorithm "peptide_auc_analysis", see Table S12 | This paper | https://github.com/VasileiosCheras/peptide_auc_analysis |
| In-house peptide barcode design Python-based algorithm "*PeptideBarcode_Generator.py*" | This paper | https://github.com/VasileiosCheras/PeptideBarcode_Generator |
| SnapGene v.4.3.11 | Snapgene | RRID:SCR_015052 |
| Fiji v2.11.0 | https://imagej.net/ | RRID:SCR_002285 |
| ImageLab 6.1 | Bio-Rad | RRID:SCR_014210 |
| MATLAB R2022a v.9.12.0.1884302 | MathWorks | RRID:SCR_001622 |
| Prism v10.1.1 software. | GraphPad Prism | RRID:SCR_002798 |
| FlowJo software v.10.8.0 | BD Biosciences | RRID:SCR_008520 |
| FlexAnalysis v.4.0 | Bruker | N/A |
| nSolver v.4.0 Analysis Software | Nanostring | SCR_003420 |
| NS_H_MIR_V3B reporter library file | Nanostring | N/A |
| Other |  |  |
| Alpha 2-4 LDplus freeze drier | Martin Christ | N/A |
| Nanodrop™ One<sup>C</sup> | ThermoFisher | Cat# ND-ONEC-W; RRID:SCR_023005 |
| Molecular Imager Gel Doc XR+ system | Bio-Rad | Cat# 1708195 |
| Nikon Eclipse Ti Microscope | Nikon | RRID:SCR_021242 |
| BD LSRFortessa II Analyzer | BD Biosciences | RRID:SCR_002159 |
| Tecan M1000Pro spectrophotometer | Tecan | N/A |
| nCounter SPRINT Profiler | Nanostring | RRID:SCR_021712 |
| rapifleX MALDI-TOF/TOF | Bruker | N/A |

## Resource Availability

### Lead contact

Further information and requests for resources and reagents should be directed to and will be fulfilled by the Lead Contact, Yaakov Benenson).

### Materials availability

Plasmids generated in this study can be made available upon request to the Lead Contact with a completed Materials Transfer Agreement.

### Data and code availability

Raw data, as well as the code generated in this study can be made available upon request.

### Peptide barcode sequence generation

An in-house generated Python script (*PeptideBarcode_Generator.py*) was used to automate the design of the peptide barcode amino acid sequences. The software is available on GitHub (https://github.com/VasileiosCheras/PeptideBarcode_Generator). The alphabet of amino acids used in the peptide barcodes was devoid of histidine, cysteine, and methionine (alphabet= ’ADEFGILNPQSTVWY’). The maximum amino acid chain length was set to 32 (max_length=32), and up to four amino acid repeats were allowed in a peptide barcodes sequence (max_repetitions=4). The total number of peptide sequences to be generated was set to 20,000 (n=20,000). The DeepMSPeptide script, a pre-compiled and pre-trained one-dimension Convolutional Neural Network, was used to calculate the detection probability of the input peptide sequences in the MS [35]. Peptides with a detection probability greater than 0.85 were considered in the design of the different miRNA sensors. All the details of the peptides used in this study, such as their amino-acid sequence, monoisotopic mass and detection probability are summarized in Table S1.

### Genetic elements of peptide-barcoded miRNA sensors

The genetic design of the peptide barcoded miRNA sensors included; (i) the constitutive bidirectional cytomegalovirus (biCMV) promoter from pMD26 [25]; (ii) the mCitrine and mCherry coding genes from pKH025 and pKH026 respectively [57]; (iii) the polyadenylation signal of the rabbit beta-globin gene (RbGpA) from pKH025 downstream of the sense expression cassette; (iv) the human growth hormone polyadenylation (hGHpA) signal from pMD26 downstream of the anti-sense expression cassette; (v) the GGS and GSS linker sequences from pEF1/His B (Invitrogen, Cat# V92220) and pSelect His-Tag (InvivoGen, Cat# psetb-chis) respectively; (vi) the 6x-histidine tag (6x-His-tag) sequence from pEZYmyc-His (Addgene Plasmid #18701) [59]; (vii) the peptide barcodes fused to the mCitrine and mCherry-based reporters as described in Table S2. All the peptide barcode, 6x-His-tag, FLAG-tag and linker sequences of the fusion reporters were codon optimized for expression in human cells using SnapGene software v4.3.11. pRA52 described by Altamura et al. served as a plasmid backbone in the design of the miRNA sensors pVC001, pVC050, pVC070, pVC071 and pVC073) [58]. A BsaI-free version of the pRA52 backbone (pVC066) was used in combination with a BsaI-free hGHpA signal (pVC077) to enable BsaI-based Golden Gate assembly for the rest of sensor plasmids [60].

### Recombinant DNA methods

All DNA plasmids were assembled using standard molecular cloning techniques, such as restriction ligation, Golden Gate cloning and Gibson assembly [60, 61]. The manufacturer protocols were followed for every kit used unless stated otherwise. All restriction enzymes used in this work were purchased from New England Biolabs (NEB, USA). Primers and oligonucleotides were ordered desalted from IDT (USA). Oligonucleotides used for oligo annealing were either ordered phosphorylated or were individually phosphorylated in-house. Synthetic gene fragments were ordered either from Twist Biosciences (USA) or IDT (USA). Phusion High Fidelity DNA Polymerase (NEB, Cat# M0530) was used in combination with a touchdown PCR protocol for DNA amplification. Restriction digestions took place at 37°C for all enzymes used, except for BstBI-based reactions where an incubation temperature of 65°C was used. Backbone plasmids were simultaneously digested using the appropriate restriction enzymes and dephosphorylated using Quick CIP (NEB, Cat# M0525L). Digested DNA products or PCR amplified products were purified using either using MinElute PCR purification kit (Qiagen, Cat# 28006) or QIAquick PCR purification kit (Qiagen, Cat# 28106). MinElute Gel purification kit (Qiagen, Cat# 28606) or QIAquick Gel Extraction kit (Qiagen, Cat# 28706) were used when gel extraction-based DNA purification was necessary. Ligation and Golden Gate Assembly reactions of plasmid backbones and inserts were implemented using T4 DNA Ligase (NEB, Cat# M0202S). Gibson assembly was implemented using a commercial Gibson Assembly Mastermix (NEB, Cat# E2611S). The protocols followed for each assembly method are described in the following sections (Ligation of DNA plasmids, Golden Gate Assembly, Gibson Assembly). Typically, 6 μL of the assembled products were transformed into in-house prepared chemically competent *Escherichia coli* TOP10 cells (ThermoFisher, Cat# C404010). The cells were plated on LB Agar plates supplemented with appropriate antibiotics (ampicillin 100 µg/mL (Sigma Aldrich, Cat# A9518) or kanamycin 50 µg/mL (Sigma Aldrich, Cat# K4000)). Identification of bacterial clones carrying the plasmids of interest, was either implemented via restriction digestion using the appropriate enzymes to generate a unique fragment pattern or by performing PCR directly on bacterial colonies with Quick-Load *Taq* 2x Master Mix (NEB, Cat# M0271) and appropriate amplification primers pairs. Selected single clones were expanded in Difco LB broth, Miller (BD, Cat# 244610) supplemented with appropriate antibiotics. DNA plasmid isolation took place using ZR Plasmid Miniprep - Classic (Zymo Research, Cat# 4016). All the plasmids were verified using either Sanger sequencing provided by Microsynth AG (Switzerland) and Eurofins Genomics GmbH (Germany) or whole-plasmid sequencing provided by Plasmidsaurus (USA). PureYield™ Plasmid Midiprep System (Promega, Cat# A2495) was used to isolate adequate DNA amounts from 100 mL of liquid bacterial cultures for mammalian cell transfections. Endotoxins were removed from the isolated DNA using the Norgen Endotoxin Removal Kit Midi (Norgen Biotek, Cat# 52200). The amount of isolated DNA was determined using agarose gel electrophoresis, as described below. Primers and DNA fragments used for plasmid cloning, are listed in Table S3 , S4 and S5. The cloning procedures of fully assembled plasmids can be found in Table S6. assembled plasmids can be found in Table S7.

#### Annealing and phosphorylation of oligonucleotides

For the annealing reaction 2.5 μL of each oligonucleotide (100 μΜ) were combined with 1 μL of TE buffer (Invitrogen, Cat# 8019005) supplemented with 500 mM NaCl (Sigma-Aldrich, Cat# S3014) and 7 μL of TE buffer. The reaction was incubated in a thermocycler at 95°C for 3 minutes (min) , followed by a decrease of 1.5°C every min for the next 57 min. Phosphorylation of the annealed products was performed by adding together 6 µL of the annealing reaction, 5 µL 10xPNK buffer (NEB, Cat# B02015), 5 µL ATP (10 mM) (NEB, Cat# P07565), 1.5 µL of T4 PNK (10 U/µL) (NEB, Cat# M0201S) and 32.5 µL RNase/DNase-free water (Invitrogen, Cat# 10977-049). The reaction was incubated at 37°C for 40 min, followed by 20 min incubation at 65°C for heat inactivation. The annealing reaction was first diluted 100-fold using RNase/DNase-free water before it was used in downstream ligation reactions.

#### PCR for DNA amplification

PCR reactions took place in 50 μL reaction volume consisting of 1 ng of DNA template, 0.25 μM of forward primer, 0.25 μM of reverse primer ,0.2 mM of deoxynucleotide triphosphate dNTPs (dNTPs, Sigma-Aldrich, Cat# D7295-2ML), 5 units of Phusion High Fidelity DNA Polymerase, 1 μL of 5x Phusion HF-Buffer and filled up to 50 μL using RNase/DNase free water (Invitrogen, Cat# 10977-049). DNA was amplified from the templates of interest, using a touchdown protocol in a PCR Thermocycler [62]. The thermocycler program was consisting of the following steps: initial template denaturation 30 seconds (sec) at 98°C; 15 cycles of 10 sec at 98°C, 30 sec at 70°C-61°C with a decrease rate of 0.5°C/cycle, 50 sec at 72°C; 15 cycles of 10 sec at 98°C, 30 sec at 61°C, 50 sec at 72°C; final DNA chain elongation for 7 min at 72°C. This protocol was suitable for the amplification of 1.5 kb fragments. For longer fragments the elongation time during the cyclic steps was increased based on the rate of DNA synthesis of 30 sec/1,000 bp.

#### Ligation of DNA plasmids

In a typical ligation reaction, 20 fmol of digested backbone plasmid were used per reaction. The molar ratio of backbone plasmid per insert was 1:5 when gene fragments were used as inserts or 1:10 when annealed oligonucleotides were used as inserts. Specifically, for a ligation reaction 1 μL T4 DNA ligase (NEB, Cat# M0202) and 2 μL 10x T4 DNA Ligase Reaction Buffer (NEB, Cat# B0202A) were combined with 2-6 μL of DNA components and supplemented with RNase/DNase free water to a final volume of 20 μL. The reactions were incubated at 16°C for 16 hours (h), followed by incubation at 65°C for 10 min, for enzyme inactivation.

#### Golden Gate Assembly

A 20 μL reaction consisted of 2 μL T4 DNA ligase (NEB, Cat# M0202), 1.5 μL BsaI-HF®v2 (NEB, Cat# R3733L), 2 μL 10x T4 DNA Ligase Reaction Buffer (NEB, Cat# B0202A), 8 μL of DNA components, and 13.5 μL RNase/DNase free water. The amount of backbone plasmid used in a Golden Gate reaction was 34.4 fmol, with a molar ratio of backbone per insert of 1:2. The reactions were incubated in a thermocycler for 59 cycles of 37°C for 5 min followed by incubation at 16°C for 5 min. At the end of the cycling program, the reactions were incubated at 60°C for 5 min, for heat inactivation.

#### Gibson Assembly

A typical Gibson reaction consisted of 10 μL of Gibson Assembly Master Mix (NEB, Cat# E2611S), 0.087 pmol of backbone DNA fragment, 0.173 pmol of insert fragments, and RNase/DNase free water to fill up the reaction to 20 μL. The reaction was incubated at 50°C for 60 min. DNA fragments carried 20 base pairs (bp) appropriate overlapping overhangs instructing the correct DNA assembly.

#### DNA quantification via agarose gel electrophoresis

For plasmid quantification, about 200 ng of plasmid DNA, as determined spectrophotometrically using Nanodrop™ One^C^ (ThermoFisher, Cat# ND-ONEC-W), were linearized in a single enzyme 20 μL restriction digestion reaction, incubated at 37°C for 2h. Then, the reaction was supplemented with 4 μL 6x Purple Gel Loading Dye (NEB, Cat# B7024S) and 12 μL were loaded onto an agarose gel prepared in-house, using 1% (w/v) agarose (Sigma Aldrich, Cat# A9539-5006) in TAE buffer (AppliChem, Cat# A4686). The gel was stained using 1X SYBR™ Safe DNA Gel Stain (ThermoFisher, Cat# S33102). On the same gel, 2, 4, 6, 8, 10 and 12 μL of GeneRuler 1 kb DNA Ladder (ThermoFisher, Cat# SM0311) were loaded. The gel was imaged using a Molecular Imager Gel Doc XR+ system (BioRad, Cat# 1708195) equipped with Image Lab Software v.6.1. The increasing band intensity of the 1 kb band of the loaded GeneRuler was used to construct an intensity versus (vs.) DNA quantity reference curve thus enabling the absolute quantification of the linearized DNA loaded on the gel. Finally, the DNA amount in the digestion reaction, and subsequently the concentration of the DNA stock aliquot was determined.

### Cell culture

The cell lines used for the experiments were HEK293 (Invitrogen, Cat# 11631-017), HeLa (ATCC, Cat# CCL-2, Lot# 58930571) and HuH-7 (Health Science Research Resources Bank of the Japan Health Sciences Foundation, Cat# JCRB0403, Lot# 07152011). 0.2 μm sterile filtered Dulbecco’s Modified Eagle Medium (DMEM) medium (ThermoFisher, Cat# 41966029), supplemented with 10% (v/v) fetal bovine serum (FBS) (ThermoFisher, Cat# 10270-106) and 1% (v/v) Penicillin/Streptomycin solution (Corning, Cat# 30-002CI), was used for cell line cultivation. Cells were incubated at 37°C and 5% CO_2_ and were split every 3-4 days using 0.25% (w/v) Trypsin-ethylenediaminetetraacetic acid (EDTA) solution (ThermoFisher, Cat# 25200-072). They were propagated for at most 20 passages before being replaced by a fresh cryopreserved cell aliquot. Mycoplasma tests were performed regularly according to the PCR Mycoplasma test kit (Promokine, Cat# PK-CA91-1024) protocol with primers specific for contaminating mycoplasmas [63]. The PCR reactions were incubated in a thermocycler with the following program: 1 cycle of 7 min at 95°C, 3 min at 72°C and 2 min at 65°C; 32 cycles of 4 sec at 95°C, 8 sec at 50°C and 45 sec at 68°C. The primers used for the detection reaction were PR1843, PR1844, PR1845, PR1846, PR1847 and PR1848. In addition to this primer mix, two more primers (PR0673 & PR0674) were included for the positive control reactions. All the primer sequences used for mycoplasma detection reactions can be found in Table S8.

### Transfection

Transfections were based on the Lipofectamine 2000 transfection reagent (ThermoFisher, Cat# 11668-027) according to the suggested guidelines. For single miRNA sensor validation experiments using flow cytometry, transfections took place in 24-well plates (ThermoFisher, Cat# 142475), while for peptide barcode purification and MS experiments 100mm plates (TPP, Cat# 93100) were used. For all experiments, cells were seeded 24 h prior to transfection. In the case of HEK293 cells, seeding densities were 6.82*10^4^ cells on 24-well plates and 3*10^6^ cells on 100mm plates. HeLa and HuH-7 cells were seeded at a density of 6*10^4^ on 24-well plates or 2.64*10^6^ on 100mm plates. After transfection optimization, the HuH-7 seeding density was changed to 6.5*10^4^ on 24-well plates or 2.86*10^6^ on 100mm plates. The cell seeding medium consisted of DMEM supplemented with 10% (v/v) FBS and 1% (v/v) Penicillin/Streptomycin solution, and the seeding volume was 500 μL (24-well plate) or 12 mL (100mm plate). The seeded cells were incubated at 37°C and 5% CO_2_ and reached approximately 80% confluency by the time of transfection.

For single transfected miRNA sensor plasmids in a 24-well plate format, 180 ng of DNA were used for all three cell lines. After transfection optimization, 275 ng and 400ng of sensor DNA were transfected into HeLa and HuH-7 cells respectively. For flow cytometry signal compensation transfections in a 24-well plate format, 180 ng of the plasmid pKH025 (Ef1a-mCitrine) or pKH026 (Ef1a-mCherry) were transfected. After transfection optimization, the compensation control samples for HeLa and HuH-7 samples were supplemented with 95 ng and 220 ng of junk DNA (pBH265 [64]). In transfection samples without any sensor or compensation-relevant plasmid, 180 ng of junk DNA were used for all the three cell lines. After transfection optimization, 275 ng and 400ng of junk DNA were transfected into HeLa and HuH-7 cells, respectively.

For transfections of miRNA or mock sensor plasmids in a 100mm plate, 7.92 μg of total DNA were used for all three cell lines. After transfection optimization, 12.1 μg and 17.6 μg of total DNA were transfected into HeLa and HuH-7 cells respectively. In the case of sensor library transfection, equal amounts of each sensor plasmid were mixed, to achieve the aforementioned total DNA amount. In transfection samples without any sensor plasmids, 7.92 μg of junk DNA were used for all the three cell lines. After transfection optimization, 12.1 μg and 17.6 μg of junk DNA were transfected into HeLa and HuH-7 cells respectively.

For miRNA sensor titration experiments in a 24-well plate format, increasing amounts (0, 0.09, 0.19, 0.38, 0.75, 1.5 and 3 pmol) of siRNA FF4 were used per transfection. The transfection samples were topped to 3 pmol of total siRNA using miRIDIAN negative control #2 (Dharmacon, Cat# CN-002000-01-05). For miRNA sensor titration experiments in a 100mm plate format, increasing amounts (0, 2.25, 4.5, 9, 18, 36 and 72 pmol) of siRNA FF4 were used per transfection. The transfection samples were topped to 72 pmol of total siRNA using miRIDIAN negative control #2 (Dharmacon, Cat# CN-002000-01-05). The sequences of all the siRNAs used in this study can be found in Table S9.

For each transfection, appropriate amounts of siRNAs and/or plasmids were mixed in Opti-minimal essential medium (Opti-MEM, ThermoFisher, Cat# 31985-062) to reach a total volume of 50μL (24-well plate) or 1.2 mL (100mm plate) (DNA Opti-MEM mix). A Lipofectamine-Opti-MEM solution was freshly prepared by mixing 2 μL of Lipofectamine 2000 per 1 μg of DNA or DNA-siRNA in the transfection mix and filling it up to 50μL (24-well plate) or 1.2 mL (100mm plate) with Opti-MEM. The Opti-MEM-Lipofectamine mix was incubated at room temperature for 5 min before mixing it with the DNA -Opti-MEM mix. The final transfection mix was gently mixed and incubated at room temperature for 20 min, before being dropwise added to the cells.

### Flow cytometry

Flow cytometry measurements of transfected cells took place between 40-48 h post-transfection. The cells were prepared by exchanging the spent culture medium with 150 μL (24-well plate) or 3 mL (100mm plate) of Accutase (ThermoFisher, Cat# A11105-01). They cells were incubated for 10 min at room temperature, and the cell suspension was subsequently transferred to micro-dilution tubes (Life Systems Design, Cat# 02-1412-0000). In the case of 100mm plates, the plates were washed twice with 2 mL of PBS 1x, pH = 7.4 (ThermoFisher, Cat# 10010023) to collect any remaining cells. Flow cytometry analysis was performed on a BD LSR Fortessa II Cell Analyzer (BD Biosciences). Sphero Rainbow Calibration Particles 8-peak beads (Spherotech, Cat# PCP-30-5A) were used for calibration prior to analysis. The voltage was adjusted so that the mean signal value of the strongest peak of the 8-peak beads remained constant across different experiments. The wavelengths of the excitation lasers (Ex) and emission filters (Em) used for respective fluorescent protein measurements were as follows: mCitrine (Ex: 488 nm, Em: longpass filter 505 nm followed by bandpass filter 530/11 nm), and mCherry (Ex: 561 nm, Em: longpass filter 600 nm followed by bandpass filter 610/20 nm).

### Microscopy

Fluorescence microscopy was used to examine the expression of the different fluorescent reporters 40-48 h post-transfection. A Nikon Eclipse Ti microscope equipped with a mechanized stage, a temperature control chamber at 37°C, and a Hammamatsu ORCA R2 camera, was used for image acquisition. The excitation light was generated by a Nikon IntensiLight C-HGFI fiber illuminator and filtered through a set of optimized Semrock filter cubes. Each cube comprised an excitation bandpass filter, an emission bandpass filter, and a specific dichroic filter for each individual fluorescence reporter. Specifically, the YFP HC filter (HC 500/24, HC 542/27, BS 520) and TxRed HC filter (HC 624/40, HC 562/40, BS 593) were used for mCitrine and mCherry, respectively. A 10x magnification lens was used in every image acquisition. The exposure time and look-up tables (LUT) for all experiments are indicated in the respective figure legends. The Fiji software v2.11.0 was used for image processing.

### Protein and peptide barcode purification

#### Cell isolation for protein purification

Peptide-barcoded reporter proteins were isolated from cells transiently transfected on 100mm plates. The single-cell suspension of each sample was pelleted by centrifugation (250 g, 4°C, 9 min). Afterwards, the supernatant was discarded, and cells were resuspended in 6 mL cold PBS 1x, pH = 7.4. This process was repeated twice. During the last washing step, the cell concentration was determined by quadruplicate measurements on a TC-10 automatic cell counter (BioRad, USA). 2-10*10^6^ cells were transferred in a fresh tube and pelleted via centrifugation for further processing.

#### Nickel-nitrilotriacetic-(Ni-NTA) based protein purification

For Ni-NTA-based affinity purification, the isolated cells were resuspended in 500 μL of cell lysis solution consisting of 0.08% (v/v) Tween 20 (Sigma-Aldrich, Cat# P9416-100ML), 10 mM imidazole (Sigma-Aldrich, Cat# 56749-50G), 250 mM NaCl (Sigma-Aldrich, Cat# S3014-1KG) in deionized water and regulated at pH=8. The resuspended cells were transferred to 1.5 mL protein LoBind tubes (Eppendorf, Cat# 0030108116), and cell lysis took place by incubation at 4°C for 30 min on a rotating shaker. For experiments performed under optimized conditions, the Tween 20 reagent was exchanged with 0.06% (v/v) Triton X-100 (AppliChem, Cat# A1388.0500). The cell lysates were centrifuged (14,000 g, 4°C, 10 min) and the supernatants were transferred to fresh protein LoBind tubes kept on ice. Ni-NTA agarose (20 μL per sample, QIAGEN, Cat# A1388.0500) was equilibrated in 5 volumes of Ni-NTA binding solution (10 mM imidazole and 250 mM NaCl in deionized water regulated at pH = 8), by three rounds of centrifugation (500 g, 4°C, 3 min) and supernatant removal in between. The pelleted agarose was resuspended in Ni-NTA binding solution (50 μL per sample) and distributed to the different cell lysates. The samples were incubated at 4°C for 150 min on a rotating shaker for Ni-NTA-based protein capture. The agarose was then pelleted via centrifugation, resuspended in 500μL Ni-NTA wash buffer (20 mM imidazole and 250 mM NaCl in deionized water regulated at pH = 8), and incubated for 5 min at room temperature to remove non-specifically bound proteins from the resin. This process was repeated twice.

#### Anti-FLAG-based protein purification

For anti-FLAG immunoaffinity purification, the DYKDDDDK Fab-Trap Agarose Kit (ChromoTek, Cat# ffak-20) was used. Specifically, the isolated cells were resuspended in 200 μL of cell lysis solution (ChromoTek, Cat# lysbuf-30), transferred to 1.5 mL protein LoBind tubes, and incubated at 4°C for 30 min on a rotating shaker. 25 μL of anti-FLAG agarose slurry were distributed in separate protein LoBind tubes. The resins were equilibrated by three rounds of buffer exchange using 300 μL dilution buffer (ChromoTek, Cat# dbuf-10), centrifugation (1,000 g,4°C, 3 min), and supernatant removal. After complete cell lysis, the lysates were centrifuged (15,000 g, 4°C, 10 min) and the supernatants were distributed onto the equilibrated anti-FLAG agarose. The lysate-bead suspensions were incubated at 4°C for 150 min on a rotating shaker for immunoaffinity-based protein capture. The protein-loaded agarose was then pelleted (1,000 g, 4 °C, 3 min) and the supernatant was removed. Any residual lysate from the resins was removed by washing three times with 500 μL of wash buffer (ChromoTek, Cat# wbuf-10), centrifugation (1,000 g, 4°C, 3 min), and removal of the supernatant.

#### On-resin peptide cleavage

Protein-loaded resins (either Ni-NTA or anti-FLAG agarose) were washed twice with 500 μL Enterokinase reaction buffer (50 mM NaCl, 20 mM Tris/Cl (ThermoFisher, Cat# 15568-025), 20mM CaCl_2_ (Merck, Cat# 1.02382.0500) in deionized water regulated at pH=8), of centrifugation (1,000 g, 3 min), and removal of the supernatant. The pelleted resins were resuspended in 20 μL of Enterokinase reaction buffer supplemented with 16 U of Enterokinase light chain (NEB, Cat# P8070L) by carefully tapping each tube. On-bead proteolytic cleavage of peptide-barcoded proteins was performed for 22 h at 25°C, 1,200 rpm using a ThermoMixer (Eppendorf, USA).

In the experiment depicted in Figure 8, the enterokinase was removed by adding 20 μL of trypsin-inhibitor agarose (Sigma-Aldrich, Cat# T0637-5ML) per sample. The resin was previously equilibrated by three washing steps involving the addition of at least 5× excess of enterokinase reaction buffer, centrifugation (1,000 g, 3 min), and removal of the supernatant. In the end, the resin was resuspended in enterokinase reaction buffer, distributed to the samples, and incubated for 1 h at 25°C, 1,200 rpm.

In preparation for isolating the cleaved peptides from the enterokinase/resin mix, empty Micro Bio-Spin^TM^ chromatography columns (BioRad, Cat# 7326204) were conditioned once with 100 μL of MS-grade water (Merck, Cat# 1.15333.2500), and placed into new protein LoBind 1.5 mL tubes. 30 μL of MS-grade water were added to each enterokinase/resin mix, and the whole sample was transferred to the conditioned columns. The loaded chromatography columns were centrifuged (1,000 g, 3 min), and the eluates containing the cleaved peptides were kept on ice. The resin on the chromatography column was washed twice with 75 μL of MS-grade water. The eluates of all centrifugation steps for a specific sample were combined into a protein LoBind 0.5 mL tube (Eppendorf, Cat# 0030108094). The peptide isolates were frozen in liquid nitrogen, lyophilized using an Alpha 2-4 LDplus freeze drier (Martin Christ, Germany), and stored at -20°C for further processing.

### Fluorescence Spectroscopy

Fluorescence spectroscopy was used to evaluate the binding capacities of Ni-NTA and anti-FLAG resins, by detecting the mCitrine- and mCherry-based reporters in different lysates before and after protein purification. Equal amounts of cells, transfected with mock sensors, were lysed using either 600 μL Ni-NTA lysis buffer or anti-FLAG lysis buffer. 50 μL of each lysate were diluted by adding 50 μL of the corresponding lysis buffer and kept for downstream analysis. 500 μL of each lysate were mixed with 50 μL of Ni-NTA or anti-FLAG resins. After protein capturing, the resins were pelleted, and ∼550 μL of each supernatant lysate were isolated. 100 μL of the lysate before and after capturing were transferred in a flat bottom black polystyrene 96-well plate (Merck, Cat# M0562) and analyzed using a Tecan M1000Pro spectrophotometer (Tecan, Switzerland). mCitrine fluorescence was detected using 488 nm excitation/ 529 nm emission wavelength, while mCherry fluorescence was detected using 582 nm excitation/ 613 nm emission wavelength. Both the excitation and emission wavelengths had a bandwidth of 5 nm, and the number of flashes was set to 50 at 400 Hz frequency. The gain value was set to 98 and 143 for mCitrine and mCherry measurements, respectively.

### Mass spectrometry

#### Sample preparation

Synthetic peptides were ordered either from PepScan (Netherlands) or GenScript (USA), and diluted in 0.1% (v/v) MS-Grade trifluoroacetic acid (TFA, Merck, Cat# 1.59014.2500). Lyophilized peptides isolated from cell lysates were resuspended in 15 μL 0.1% (v/v) TFA prior to MALDI-TOF analysis. MALDI-TOF matrix solution was prepared by dissolving α-cyano-4-hydroxycinnamic acid powder (Sigma-Aldrich, Cat# 70990) in TA30, TA50 or TA70 buffer (30 or 50 or 70 % MS-Grade Acetonitrile with 0.1 % (v/v) TFA (Merck, Cat# 1.59014.2500), 70 or 50 or 30% 0.1% (v/v) TFA) to achieve a final concentration of 10-18 mg/mL. MS sample preparation took place by mixing 5-8 μL of a peptide sample with appropriate amounts of a synthetic spike-in peptide diluted in 0.1% (v/v) TFA. The peptide mixture was mixed 1:1 with freshly prepared matrix solution and briefly vortexed. 1-2 μL of the resulting solution were spotted on a freshly cleaned MTP 384 ground steel target plate (Bruker, Cat# 8280784). The samples were left to crystallize at room temperature under a chemical hood. The sample preparation followed across different experiments are summarized in Table S10.

#### MALDI-TOF analysis

A rapifleX MALDI-TOF/TOF system (Bruker, USA) was used to acquire MS1 spectra of the crystallized samples. The sample acquisition was implemented using reflector positive ion mode, 600 – 3200 m/z detection range and optimized detector gain per experiment (x3.14 or x5.17). Tο allow meaningful data comparisons, experiments with identical gains were used for comparisons. The laser power was adjusted for each experiment (30%-70%), and the laser frequency was set to 10 kHz. Each spot was sampled randomly, accumulating 40,000 shots in 250 shot steps. The cumulative spectrum of each spot was saved in a container format (.fid) and converted to a .txt file using the FlexAnalysis v.4.0 software (Bruker, USA). The MS acquisition parameters followed across different experiments are summarized in Table S9.

#### miRNA abundance profiling

To estimate the miRNA abundance in HEK293, HeLa and HuH-7 cells, duplicate samples, each consisting of 5*10^6^ cells, were isolated per cell line. Each sample was washed with 6 mL of PBS 1x, pH = 7.4 and centrifuged (250 g, 9 min). The supernatant was discarded, and cells were resuspended in 3 mL of PBS 1x, pH = 7.4. 1 mL of the cell solution (1.67*10^6^ cells) was further used for total RNA isolation using the miRVana miRNA Isolation kit (ThermoFisher, Cat# AM1560). The quantity and quality of the isolated RNA were evaluated spectrophotometrically using Nanodrop™ One^C^ (ThermoFisher, Cat# ND-ONEC-W). Each sample was diluted to 33 ng/μL with RNase/DNase free water (Invitrogen, Cat# 10977-049). The miRNA abundance in each sample was evaluated via the nCounter miRNA expression assay (Nanostring, USA) using 100 ng of the total RNA solution per sample. Human miRNA-tag (Nanostring, Cat# Hu-MIRTAG-12) ligation and human miRNA CodeSet (Nanostring, Cat# CSO-MIR3-12) hybridization steps were implemented according to the manufacturer’s protocol. 20 μL of the hybridized product per sample were injected into a nCounter cartridge (Nanostring, Cat# SPRINT-CAR-1.0) and analyzed using the nCounter SPRINT Profiler instrument (Nanostring, USA). The reporter code count (.RCC) result files were converted to raw detected miRNA probe counts per sample using the nSolver v.4.0 Analysis Software (Nanostring, USA) and the NS_H_MIR_V3B reporter library file.

### Data analysis

#### Flow cytometry data analysis

FlowJo software v10.8.0 (BD Biosciences) was used for flow cytometry data analysis. The following steps were followed to analyze the acquired data. Firstly, live cells were gated based on forward scatter area (FSC-A) versus side scatter area (SSC-A). The live cell population was plotted on FSC-A versus forward scatter height (FSC-H), and single cells were gated. The fluorescence values of single cells were corrected using a compensation matrix to account for spectral overlap between fluorescent proteins (Table S11). The matrix was based on cells transfected with constitutively expressed mCitrine (pKH025) or mCherry (pKH026). To distinguish mCitrine and/or mCherry-expressing cell populations, fluorescent values of single cells transfected with junk DNA were plotted on histograms for each fluorescent channel and cells were gated, such that 99.9% of cells fell outside of the selected gate [Fig. S12]. This gating is applied throughout the single-cell population of all samples to determine each positive single-cell population.

The frequency of positive cells and the mean value of fluorescence for every fluorescence channel are summarized for each sample in a statistics table generated via FlowJo. The product of these two values provides the absolute fluorescence intensity.

Dividing the absolute intensity of the peptide-barcoded mCitrine reporter (activity reporter) by that of the peptide-barcoded mCherry reporter (normalization reporter) defines the flow cytometry-based output of a miRNA or mock sensor. The following equation represents the calculation:

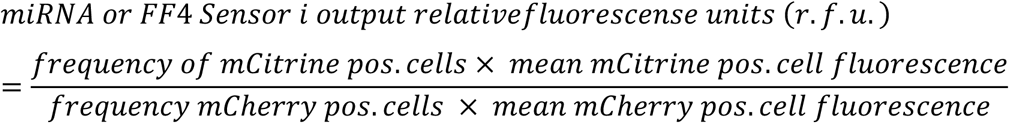

The activity of a miRNA measured via flow cytometry is determined by comparing the output of a sensor in the presence or absence of miRNA regulation, as previously described [25]. To this end, the output of a sensor carrying endogenous miRNA targets is divided by the output of a mock sensor harboring FF4 targets. To convert this ratio into a miRNA activity value, the quotient is minus log transformed.

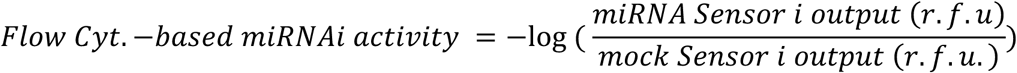

To measure the repression strength of siRNA at a given concentration during a sensor titration experiment, the sensor output is compared to that in the absence (0 nM) of the siRNA. The ratio thereof is minus log-transformed to calculate the siRNA activity.

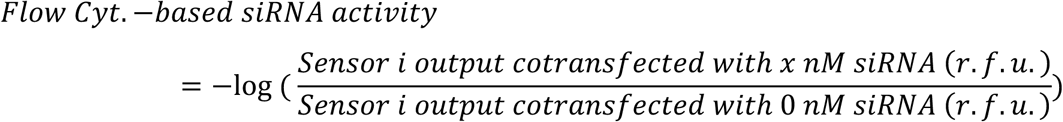

#### Fluorescence spectroscopy data analysis

The absolute fluorescence values of peptide-barcoded mCitrine and mCherry reporters present in the lysate before and after cell lysis were acquired via a spectrophotometer. These values were used to estimate the percentage of reporters captured by the Ni-NTA or anti-FLAG resins. The following equation was used to calculate percentage of captured fluorescence:

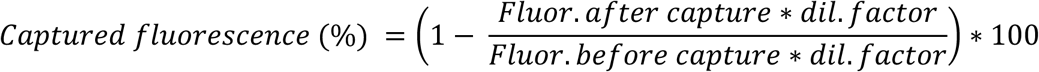

#### MS data analysis

A mass spectrum processing software tool, comprising a series of scripts written in MATLAB programming language (MATLAB R2022a v.9.12.0.1884302), was created to automate the calculation of peptide ion abundances from raw MS data. The software is available on GitHub (https://github.com/VasileiosCheras/peptide_auc_analysis). The repository contains the main function (“peptide_auc_analysis.m”) used to analyze raw spectra in .txt format, as well as all the necessary scripts in the “src” folder. A detailed user manual (“peptide_auc_analysis User Manual”), a spectra test dataset, and input parameter test files are also provided. User-provided data can be analyzed according to the steps described in the user manual.

Briefly, spectra processing proceeds as follows: (i) the user needs to provide a folder with the unprocessed (raw) spectra files in .txt format, together with three files containing information about the biological sample of origin of each spectrum, the peptide of interest expected to be found in the spectra, and a table with input parameters needed for spectra processing; (ii) given those inputs, the function stores information, such as the m/z ratio and intensity values comprising the raw spectrum, its file name, and its experimental sample of origin; (iii) the function removes m/z duplicate values from the raw spectrum; (iv) background subtraction is applied to each spectrum; (v) optionally, the background subtracted values are normalized using the Total Ion Count (TIC) method [65]; (vi) based on the expected m/z ratio of each peptide, segments extracted from the processed spectrum are generated; (vii) the isotopic peaks corresponding to the peptides of interest are identified, and their quantitative features (e.g., the area-under-the-curve (AUC), the maximum peak intensities) are extracted as a measure of their ion abundance; (viii) the output of a specific miRNA or mock sensor is defined as the relative ion abundance of its activity over normalization peptide using the equation below:

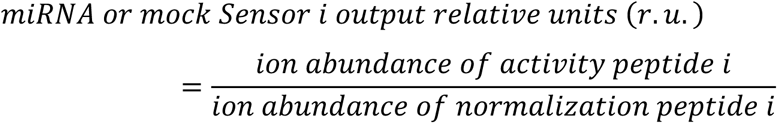

(ix) optionally, figures of processed spectra are generated, containing annotated peptide isotopic envelope peaks that visualize the peak picking algorithm output; (x) optionally, a summary of certain spectrum features, such as the absolute and relative ion abundancies sorted per experimental sample and peptide, is generated. The high-level pseudocode for the function is described in Table S12, while the input parameters used for spectra analysis can be found in Table S13. Differences in the input parameters across samples are summarized in Table S14. In this paper, the location of the maximum isotopic peak in the isotopic envelope of a peptide was not used as a criterion for peptide peak assignment. The AUC of the first two isotopic peaks of an identified peptide was used to calculate the peptide ion abundance [Fig. S13].

The activity of a miRNA measured via MS is calculated similarly to its flow cytometry-based measurement. However, in this case, the peptide barcode-defined sensor output was compared in the presence or absence of miRNA regulation. The sensor outputs were acquired from samples transfected either with a miRNA or mock sensor library. The MS-based activity of a miRNA i is calculated as follows:

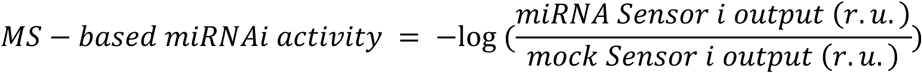

To measure the MS-based activity of siRNA at a given concentration during a titration experiment, the sensor output is compared to that in the absence (0 nM) of the siRNA. The mathematical equation describing this is shown below:

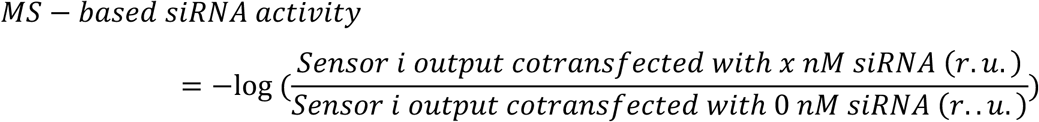

The negative logarithm of the measured activities is used in the figures to convert the decreasing miRNA sensor output to higher activity values.

Finally, to evaluate the impurities contaminating different areas of interest across the spectrum, the summed ion counts within a spectrum segment of interest spanning 3 m/z units were normalized to the TIC of the spectrum. The equation to calculate this metric is:

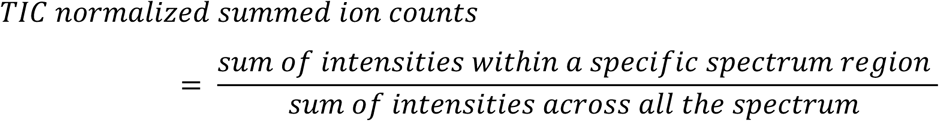

#### miRNA abundance profiling data analysis

Detected miRNA-probe raw counts, as determined from nCounter miRNA expression assay (Nanostring, USA), were background corrected using a threshold count value of 60. This value was obtained as follows: firstly, we calculated the count average and standard deviation of the negative control probes across the different experimental samples, excluding the negative control C; secondly, we added to each average two times its respective standard deviation value; next, we isolated the maximum value across the different negative controls; finally, we determined the background count threshold by doubling the isolated maximum count number. The background corrected probe count values were normalized using the top-100 method provided by the nSolver v.4.0 Analysis Software (Nanostring, USA). The normalized probe count values for each miRNA probe were converted to normalized counts per million (ncpm), and then averaged across the two biological replicate samples per cell line. The relative miRNA i subsequently log transformed to calculate the relative miRNA probe abundance detected in the different cell samples.

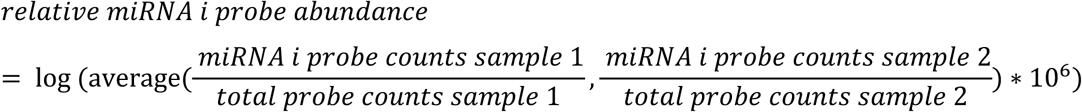

#### Statistical analysis

All the statistical analyses were performed using the built-in tools provided by Prism v10.1.1 software. Both the flow cytometry- and MS-based miRNA activity calculations stem from averaged measurements of technical replicates and /or biological replicates with distinct standard deviations. For this reason, the error propagation formula for the division of averages was used to estimate the error propagated standard deviation of the quotient. Let Av1 and Av2 be the averages of two measurement sets with standard deviations SD1 and SD2, then the standard deviation of the average quotient is calculated as:

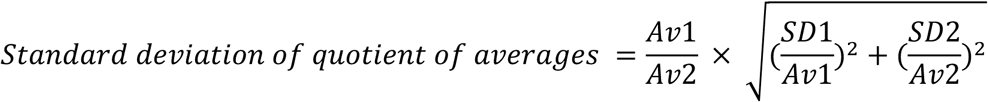

Finally, the negative logarithmic transformation used to calculate the miRNA or miRNA mimic/siRNA activities demands a second error propagation. The equation to calculate the standard deviation after the logarithmic transformation is:

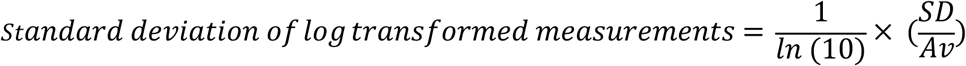

A summary of the biological replicates, technical triplicates, and statistical tests used across the different experiments can be found in Table S15.

## Supplemental information

Download 1.xlsx.Tables S1 and S2. Excel file containing the designed peptide barcode properties and information of peptide barcoded sensors.

Download 2.xlsx. Tables S3, S4, S5, S8, and S9. Excel file containing the primers, DNA fragments and siRNAs used in this paper.

Download 3.xlsx. Tables S6 and S7. Excel file containing plasmid cloning procedure and plasmid sequences.

Download 4.xlsx. Tables S6 and S7. Excel file containing plasmid cloning procedure and plasmid sequences.

Download 5.xlsx. Table S10. Excel file containing the acquisition parameters used for MALDI-TOF measurements.

Download 6.xlsx. Table S11, S12, S13, S14. Excel file containing the analysis parameters used for MALDI-TOF and Flow Cytometry measurements.

Download 7.xlsx. Table S15 Excel file containing the statistical analysis information for each paper figure.

