## Supplementary_Document_S for "Peptide Barcodes for miRNA activity assessment in mammalian cells"

### Supplementary Figures

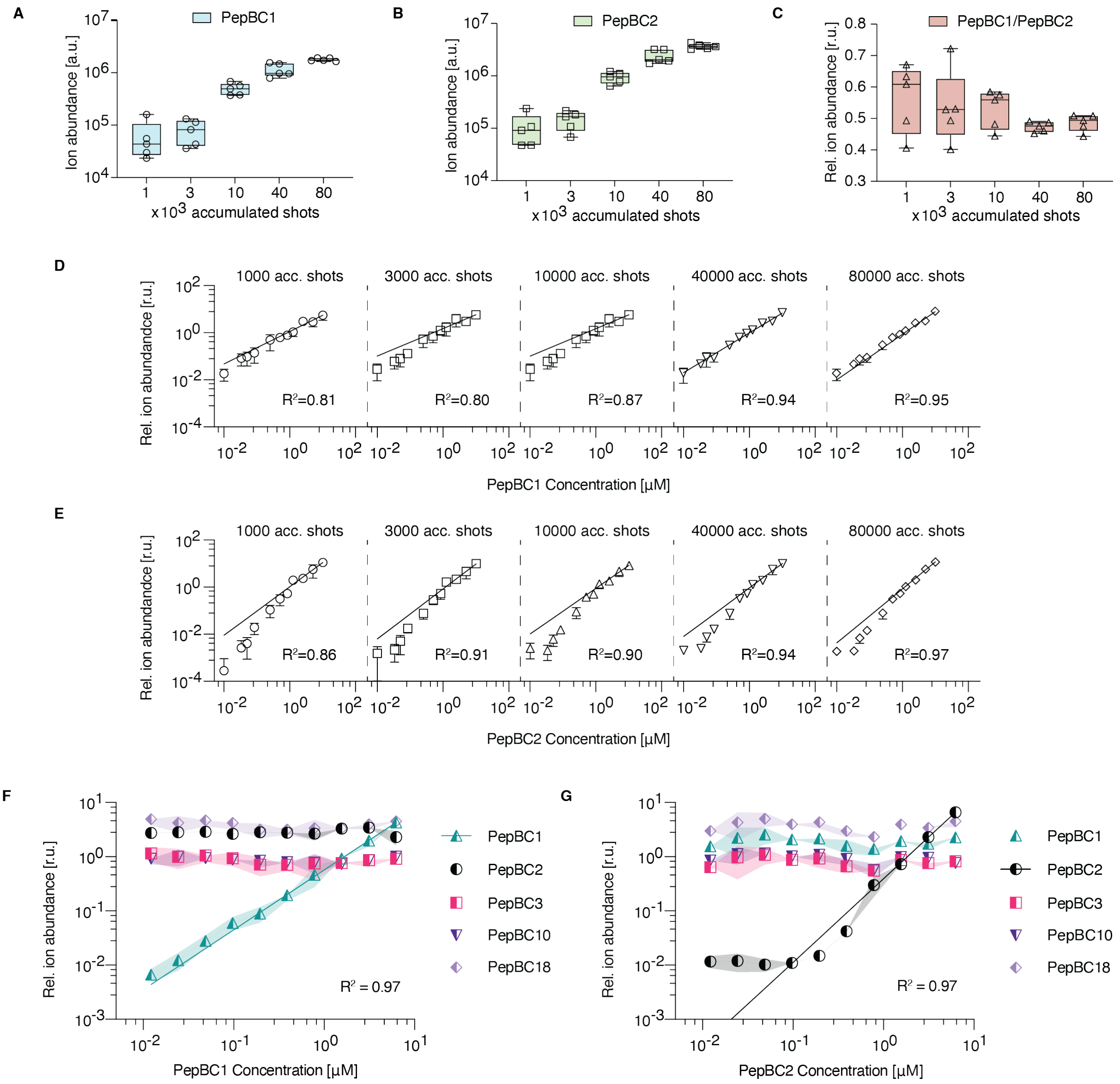


*Fig. S1*

**(A-C)** Examining the influence of increased sample coverage on the measurement of absolute and relative ion abundances of peptides in MALDI-TOF spectra. An equimolar mix of PepBC1 and PepBC2, at 1 µM each, was spotted five times on a MALDI plate. An increasing number of laser shots was used to generate spectra of increasing sample coverage. Spectrum acquisition took place starting from the lowest to the highest shot number to avoid material depletion. Three box plots are depicted, showing the ion abundance of each peptide (A-B, y-axis), as well as the relative ion abundance (C, y-axis) of the two peptides derived from spectra with an increasing number of accumulated shots (A-C, x-axis). The whiskers of the box plot represent the minimum and maximum technical replicate values acquired per condition. **(D-E)** Examining the influence of increased sample coverage on the standard curve construction for two synthetic peptide barcodes. Two standard curves were created by varying the amount of PepBC1 between 0.01 µM – 10 µM (D, x-axis) and keeping the concentration of PepBC2 steady at 1 µM, or vice versa (E, x-axis). The technical replicates were ablated with an increasing number of shots to acquire MALDI-TOF spectra with extended sample coverage, and the relative ion abundance of PepBC1 or PepBC2 was calculated for each standard curve sample (D-E, y-axis). Log-log lines were fitted to the standard curves of PepBC1 (D) and PepBC2 (E) acquired under different sample coverage settings, and their goodness of fit was evaluated via the $R^2$. **(F-G)** Peptide dilutions with concentrations between 0.0122 µM – 6.25 µM were generated for the synthetic peptide barcodes PepBC1 (F, x-axis) and PepBC2 (G, x-axis). A constant mix of PepBC2, PepBC3, PepBC10, and PepBC18, at 2.5 µM each, was included in every dilution sample of PepBC1. A constant mix of PepBC1, PepBC3, PepBC10, and PepBC18, at 2.5 µM each, was included in every dilution sample of PepBC2. PepBC19 was spiked in at 1 µM in all samples. The ion abundances of the peptides were normalized to that of the spike-in peptide within each spectrum. The relative ion abundance values across the different dilution samples were used to construct standard curves for PepBC1 (F, y-axis) and PepBC2 (G, y-axis) and to monitor the levels of the other constant peptides in each mix. Log-log lines were fitted across the different standard curves, and their goodness of fit was evaluated via the $R^2$.

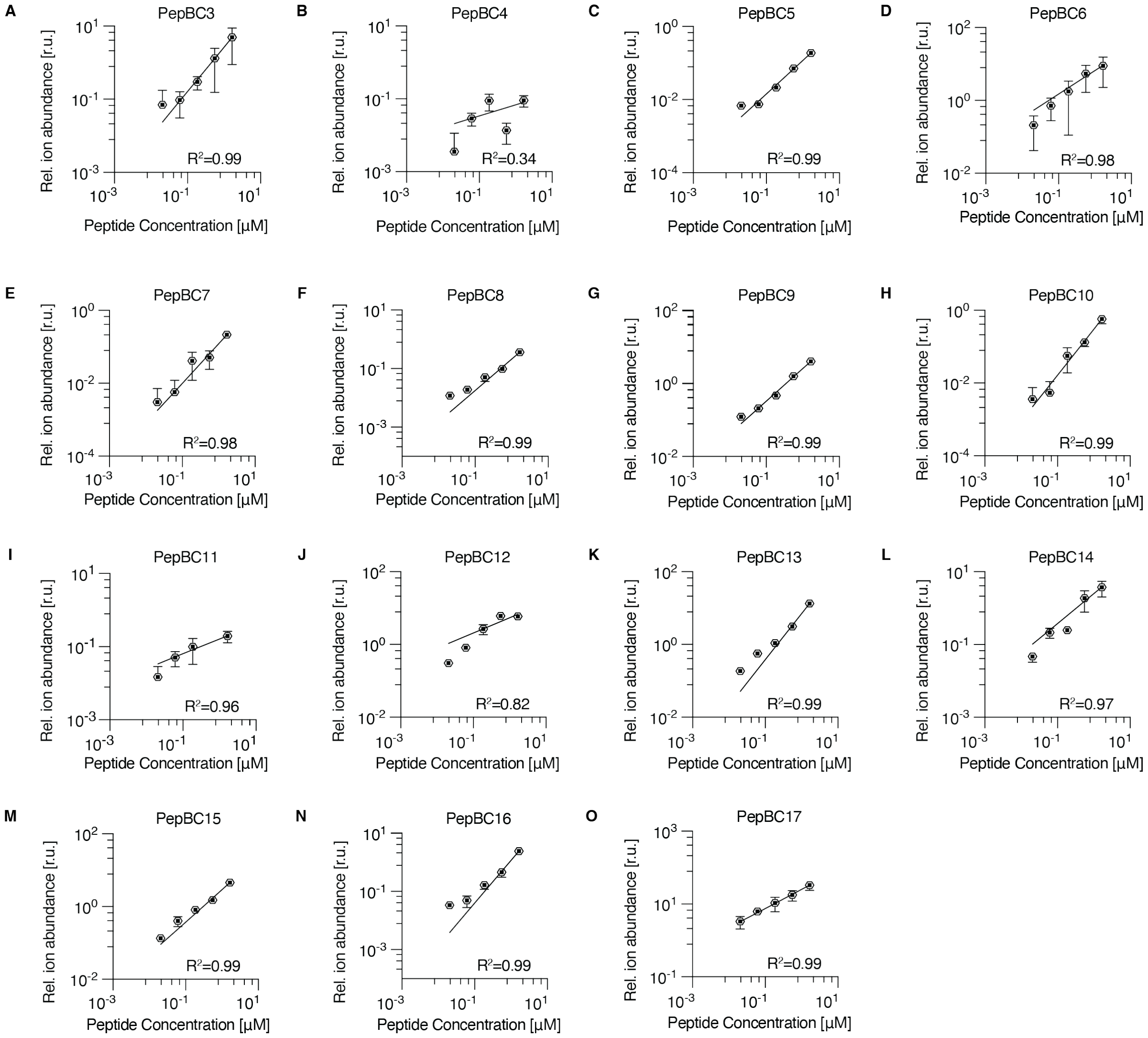


*Fig. S2*

**(A-O)** Peptide dilutions with concentrations between 0.021 µM – 1.7 µM were generated for 15 synthetic peptide barcodes (PepBC3-PepBC17). PepBC2 was spiked in at 1µM in every dilution. The ion abundances of the peptides were normalized to that of the spike-in peptide within each spectrum. The relative ion abundance values across the different dilution samples were used to construct standard curves for each peptide (A-O). Log-log lines were fitted across the different standard curves, and their goodness of fit was calculated ($R^2$). The ratio of the highest and lowest relative ion abundance observed within a standard curve was used to estimate the dynamic range of each peptide's signal in the MALDI-TOF.

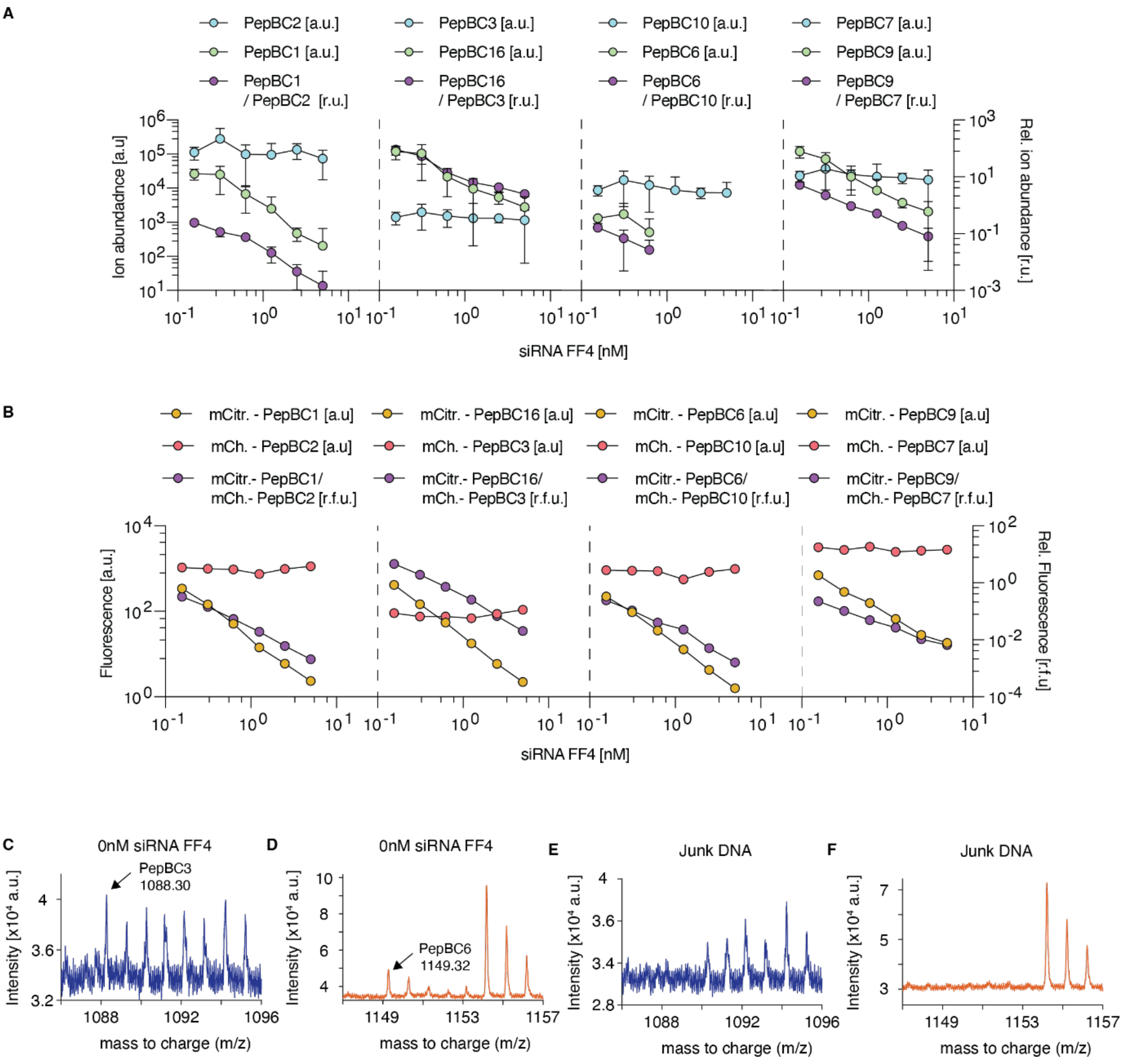


*Fig. S3.*

(**A**) Scatter plots depicting the peptide ion abundances corresponding to each sensor (left y-axis), as well as the sensor output values (right y-axis) across samples titrated with increasing siRNA FF4 concentrations (x-axis) measured via MALDI-TOF MS. Activity peptide response is depicted in green, and normalization peptide response is depicted in (blue). **(B)** Scatter plot depicting the average fluorescence values of mCitrine (yellow) and mCherry-based (red) reporters (left y-axis), as well as the average sensor output values (right y-axis, purple) across samples co-transfected with individual mock sensors and increasing siRNA FF4 concentrations (x-axis). **(C-F)** Isolated spectrum regions covering the m/z spectrum regions of PepBC3 (blue) or PepBC6 (orange). The peaks corresponding to PepBC3 or PepBC6 are denoted with an arrow and their observed m/z value. The spectra are derived from a sample transfected with the mock sensor library in the absence of siRNA mimics (0 nM) expressing the PepBC3 and PepBC6 (C,D) or a sample transfected with junk DNA, lacking any peptide barcode expression (E,F).

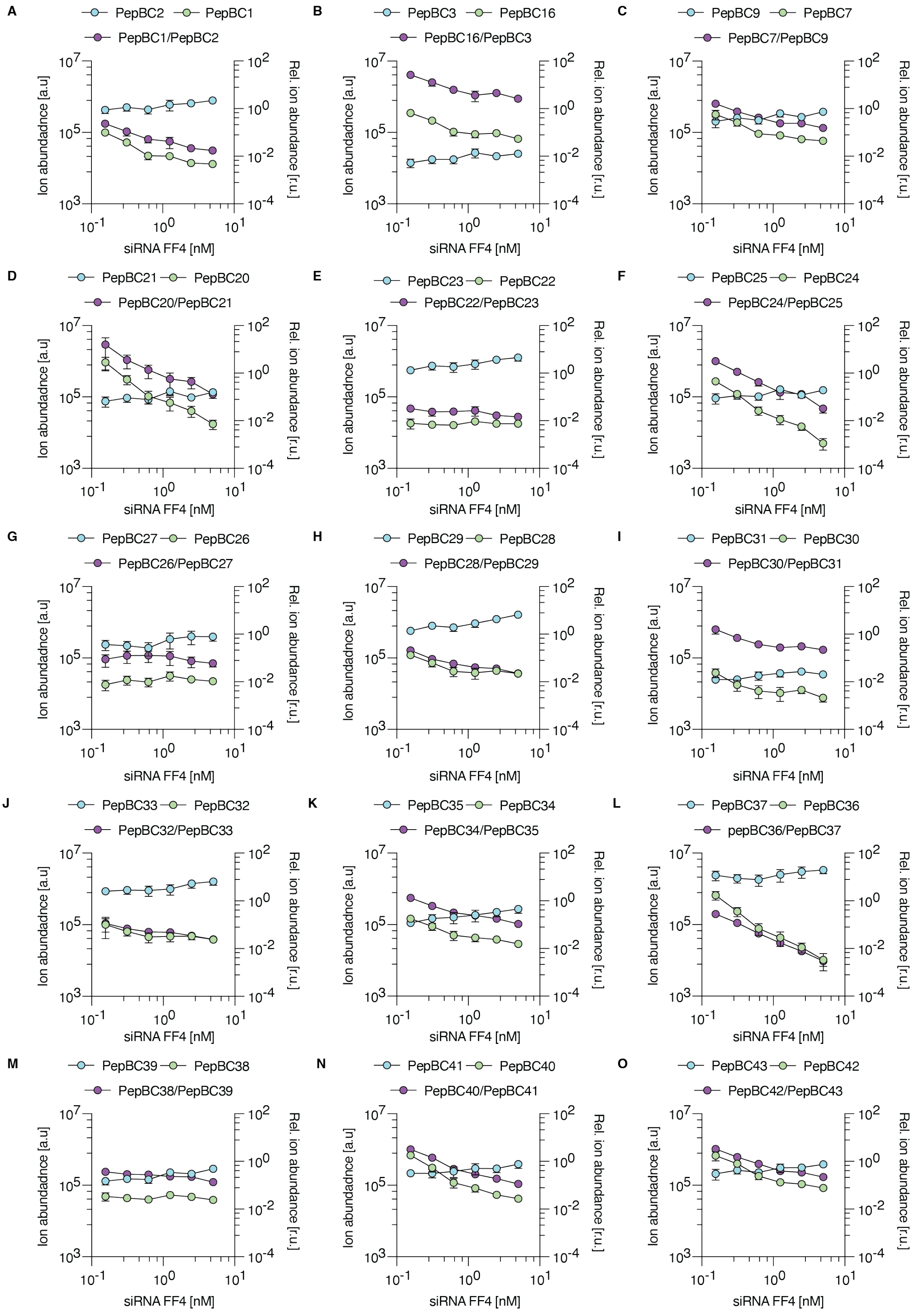

A
PepBC2
PepBC1
PepBC1/PepBC2
B
PepBC3
PepBC16
PepBC16/PepBC3
C
PepBC9
PepBC7
PepBC7/PepBC9
D
PepBC21
PepBC20
PepBC20/PepBC21
E
PepBC23
PepBC22
PepBC22/PepBC23
F
PepBC25
PepBC24
PepBC24/PepBC25
G
PepBC27
PepBC26
PepBC26/PepBC27
H
PepBC29
PepBC28
PepBC28/PepBC29
I
PepBC31
PepBC30
PepBC30/PepBC31
J
PepBC33
PepBC32
PepBC32/PepBC33
K
PepBC35
PepBC34
PepBC34/PepBC35
L
PepBC37
PepBC36
pepBC36/PepBC37
M
PepBC39
PepBC38
PepBC38/PepBC39
N
PepBC41
PepBC40
PepBC40/PepBC41
O
PepBC43
PepBC42
PepBC42/PepBC43
Ion abundadnce [a.u]
Rel. ion abundance [r.u.]
siRNA FF4 [nM]


*Fig. S4:*

**(A-O)** Scatter plots depicting the peptide ion abundances corresponding to each sensor (left y-axis), as well as the sensor output values (purple, right y-axis) across samples transfected with a 15-mock sensor library and titrated with increasing siRNA FF4 concentrations (0-5 nM) (x-axis) measured via MALDI-TOF MS. Activity peptide response is depicted in green, and normalization peptide response is depicted in blue.

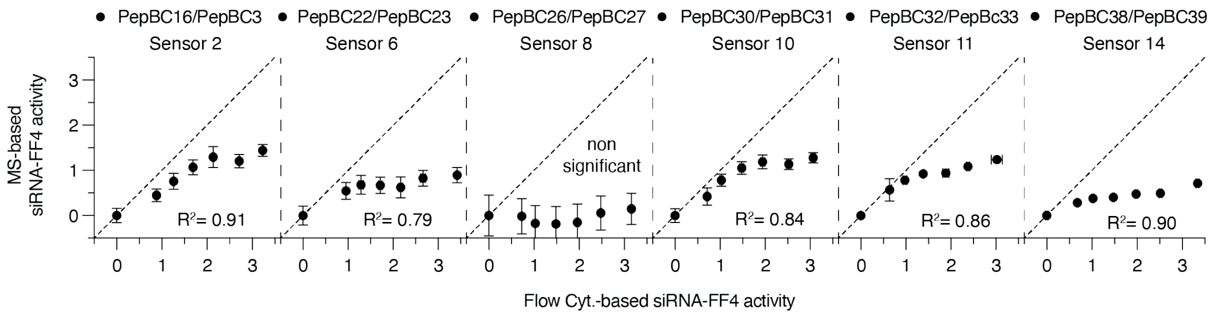


*Fig. S5:*

Scatter plots comparing the siRNA FF4 activities acquired using flow cytometry measurements of individually titrated sensors (x-axis) or MALDI-TOF MS measurements of peptides isolated from the titrated sensor library (y-axis). The sensors depicted were discarded from further use. The dashed line corresponds to the line of identity in each graph. The $R^2$ value corresponds to the result of the Pearson correlation test.

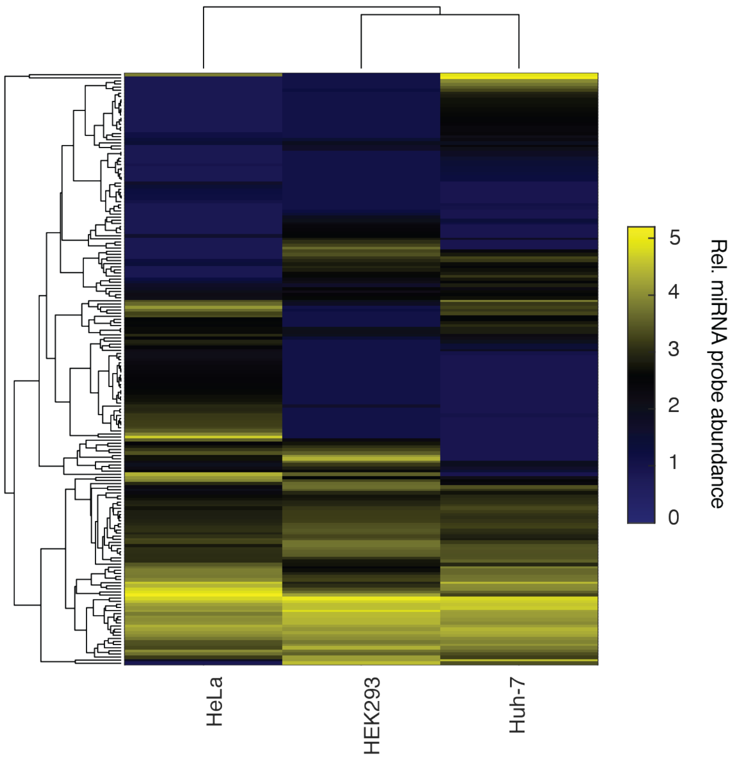


*Fig. S6:*

Clustered heatmap depicting the relative abundance of miRNA probes detected in HEK293, HeLa and HuH-7 cell samples measured via the nCounter miRNA profiling platform. miRNAs with detectable probe counts in at least one cell line are depicted.

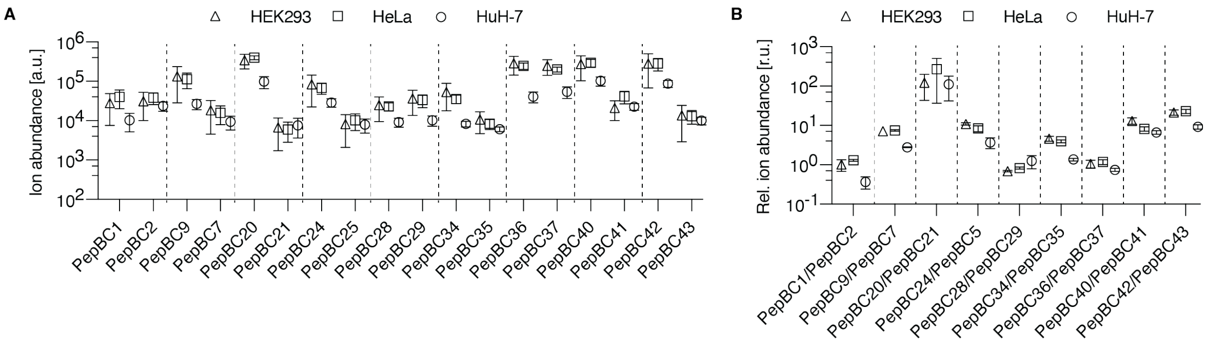


*Fig. S7:*

**(A-B)** Comparison of peptide barcode ion abundances (A, y-axis) or sensor outputs (B, y-axis) acquired from HEK293 (triangle), HeLa (square), and HuH-7 cells (circle) transfected with the mock sensor library. In panel A, activity and normalization peptide barcodes corresponding to the same sensor are separated via dashed lines.

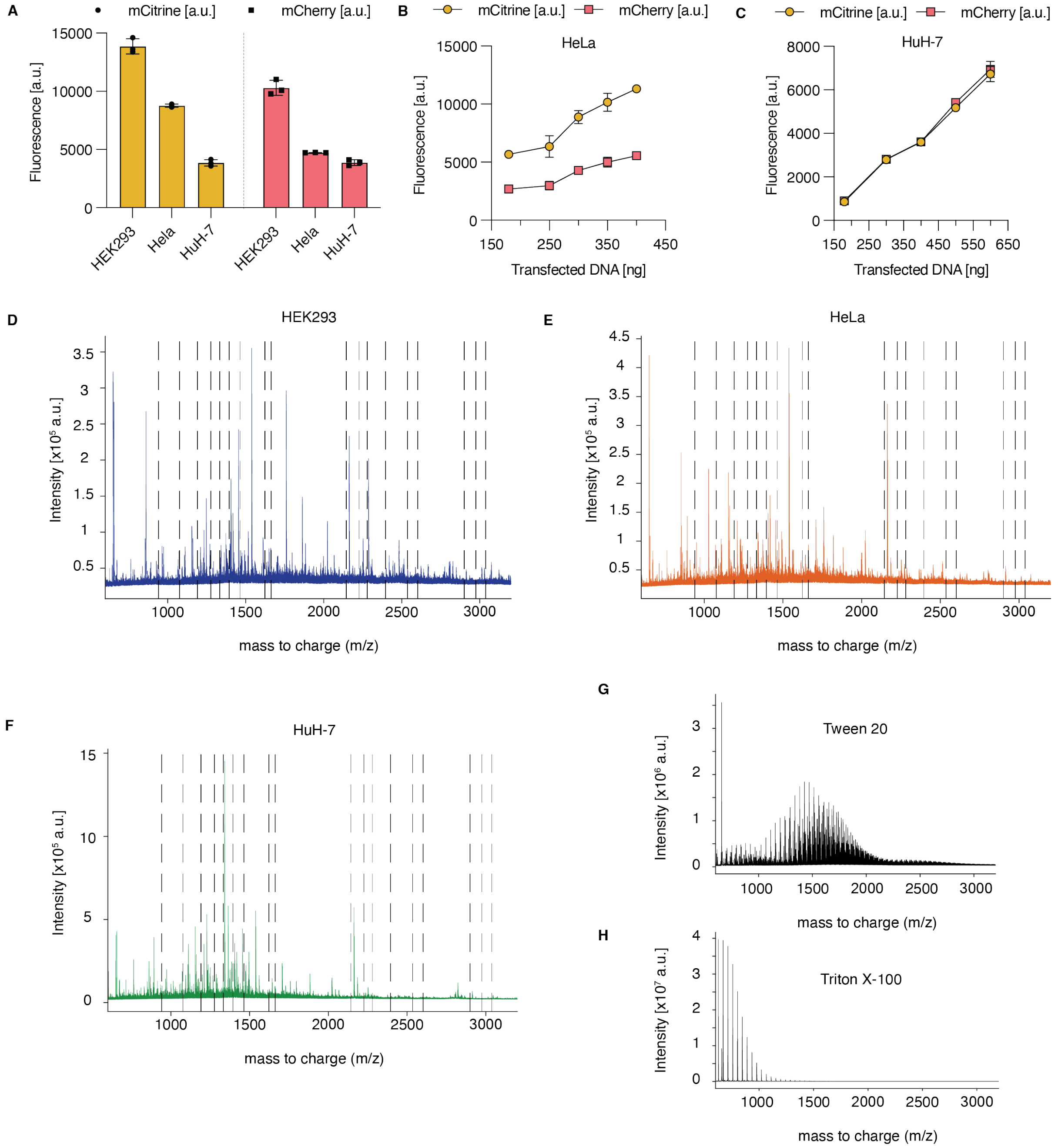


*Fig. S8:*

**(A)** Bar chart depicting the average mCitrine (yellow) and mCherry (red) fluorescence signal acquired from HEK293, HeLa, and HuH-7 cells transfected with the nine-member mock sensor library. **(B-C)** The peptide-barcoded mock sensor 13 and a four-member mock sensor library was used to evaluate the effect of increasing plasmid amount (x-axis) on the fluorescence output (y-axis) of the mCitrine- (yellow) and mCherry-based (red) reporters in HeLa (B) and HuH-7 cells (C), respectively. **(D-F)** Full spectrum of peptide isolates from HEK293 (D), HeLa (E), and HuH-7 (F) cells transfected with junk DNA. These samples lack any peptide barcode expression. Dashed vertical lines have been introduced as placeholders at the theoretical m/z positions of the peptide barcodes used in the initial miRNA sensor library. **(G-H)** Full spectrum of Tween 20 surfactant (0.08% (v/v) diluted in water) (G) and Triton X-100 surfactant (0.06% (v/v) diluted in water) (H).

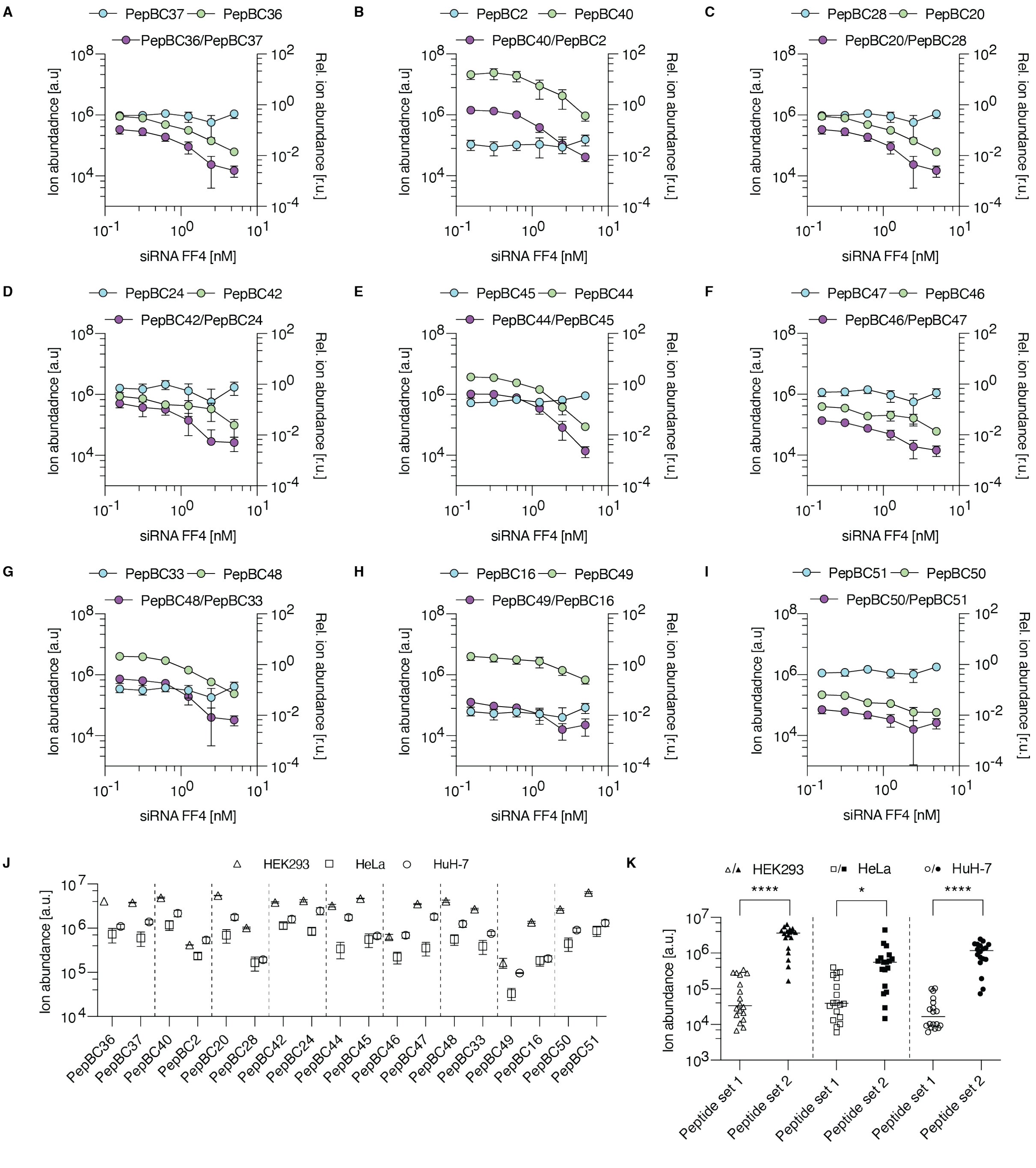


*Fig. S9:*

**(A-I)** Scatter plots depicting the peptide ion abundances corresponding to each sensor (left y-axis), as well as the sensor output values (purple, y-axis) across samples titrated with increasing siRNA FF4 concentrations (0-5 nM) (x-axis) measured via MALDI-TOF MS. Activity peptide response is depicted in green, and normalization peptide response is depicted in blue. **(J)** Comparison of peptide barcode ion abundances acquired from HEK293 (triangle), HeLa (square), and HuH-7 cells (circle) transfected with the improved mock sensor library. Activity and normalization peptide barcodes corresponding to the same sensor are separated via dashed lines. **(K)** Comparison of peptide barcode ion abundances acquired from cells transfected with mock sensors carrying either the initially selected (no fill symbols) or the optimized peptide barcode set (filled symbols). The comparison across HEK293 (triangle), HeLa (square), and HuH-7 cells (circle) is separated by dashed lines. Statistical significance of the differences observed across the peptide ion abundances between the initial and optimized conditions was calculated using an unpaired two-tailed Welch's t test (****p<0.0001, *p<0.05).

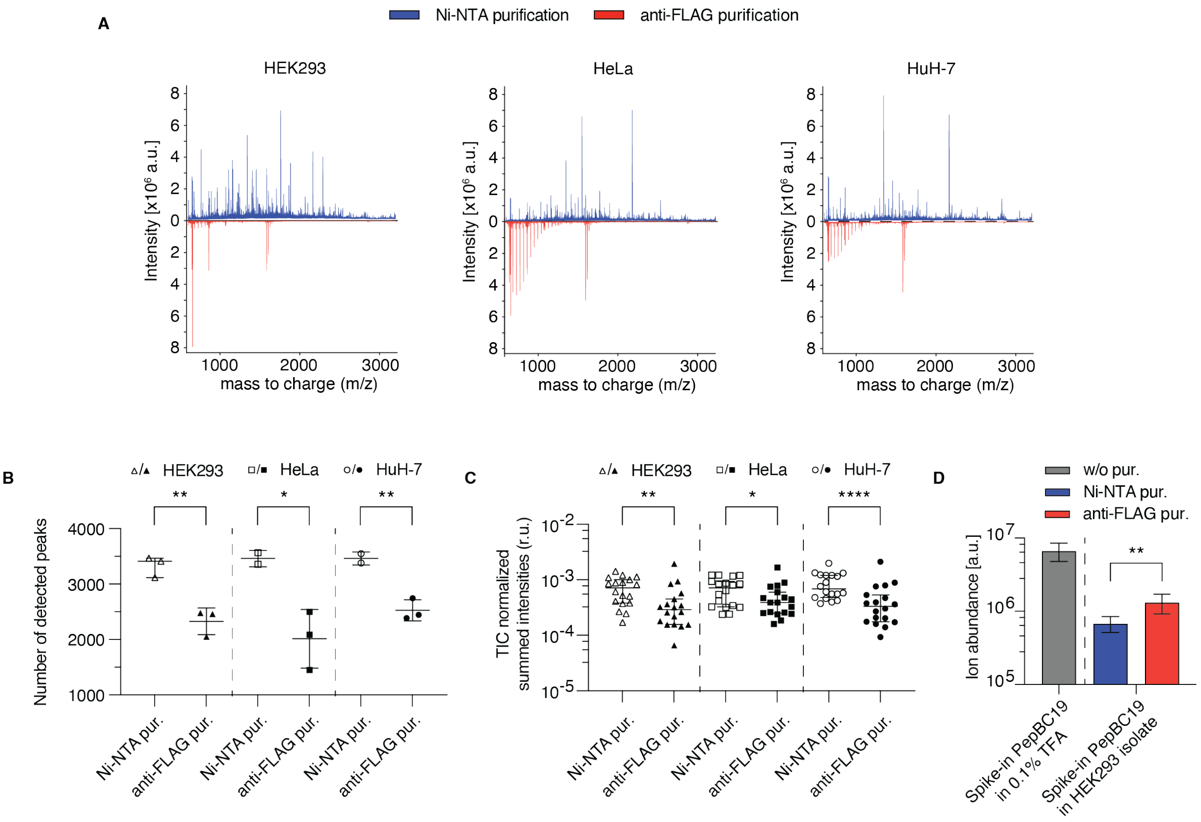


*Fig. S10:*

**(A-C)** Evaluation of the background impurities detected in the MALDI-TOF MS spectra of peptide isolates from cells transfected with junk DNA using either a Ni-NTA (no fill symbols or blue color) or an anti-FLAG protocol (filled symbols or red color). The comparison is implemented across isolates from HEK293 (triangle), HeLa (square), and HuH-7 cells (circle). The metrics compared are: (A) Full spectrum of peptide isolates from HEK293, HeLa, and HuH-7 cells transfected with junk DNA and subsequently purified via Ni-NTA or anti-FLAG purification; (B) the average number of detected peaks per spectrum; (C) the TIC-normalized summed intensities calculated for the spectrum regions covering the m/z range corresponding to the peptide barcodes of the optimized miRNA sensor set. **(E)** Bar chart depicting the ion abundance of the spike-in PepBC19 in the absence of any other peptide (grey) in the Ni-NTA or anti-FLAG peptide isolates of HEK293 cells transfected with junk DNA. Statistical significance of the differences observed across Ni-NTA- vs anti-FLAG- purified samples was calculated using an unpaired two-tailed Welch's t test (B-D) (*p < 0.05; **p < 0.01; ****p < 0.0001).

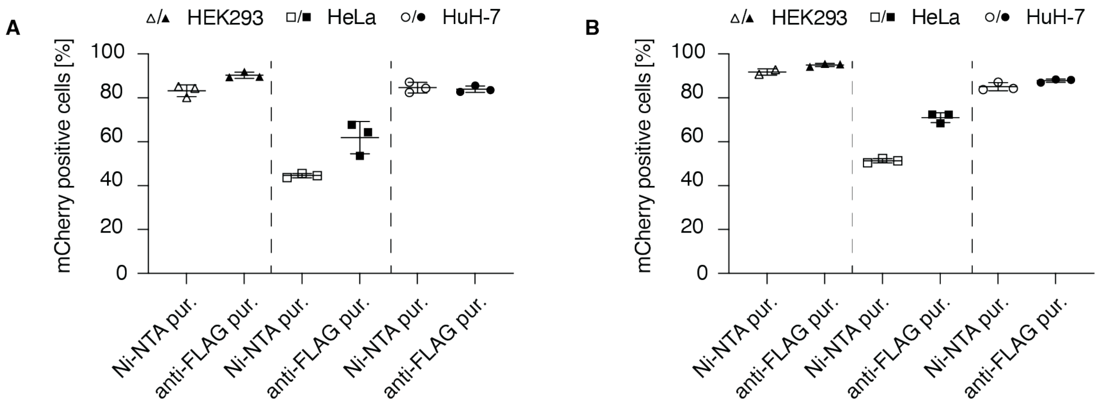


*Fig. S11:*

**(A-B)** Comparison of transfection efficiencies for cell samples purified using either Ni-NTA-based protocol (no fill symbols) or anti-FLAG based protocol (filled symbols). The transfection efficiency is evaluated using the percentage of cells that were detected positive for mCherry fluorescence, using flow cytometry. The comparison is implemented across three different cell lines, HEK293 (triangle), HeLa (square), and HuH-7 cells (circle). Samples transfected with the optimized mock sensor library are depicted in A, while samples transfected with the optimized miRNA sensor library are depicted in B.

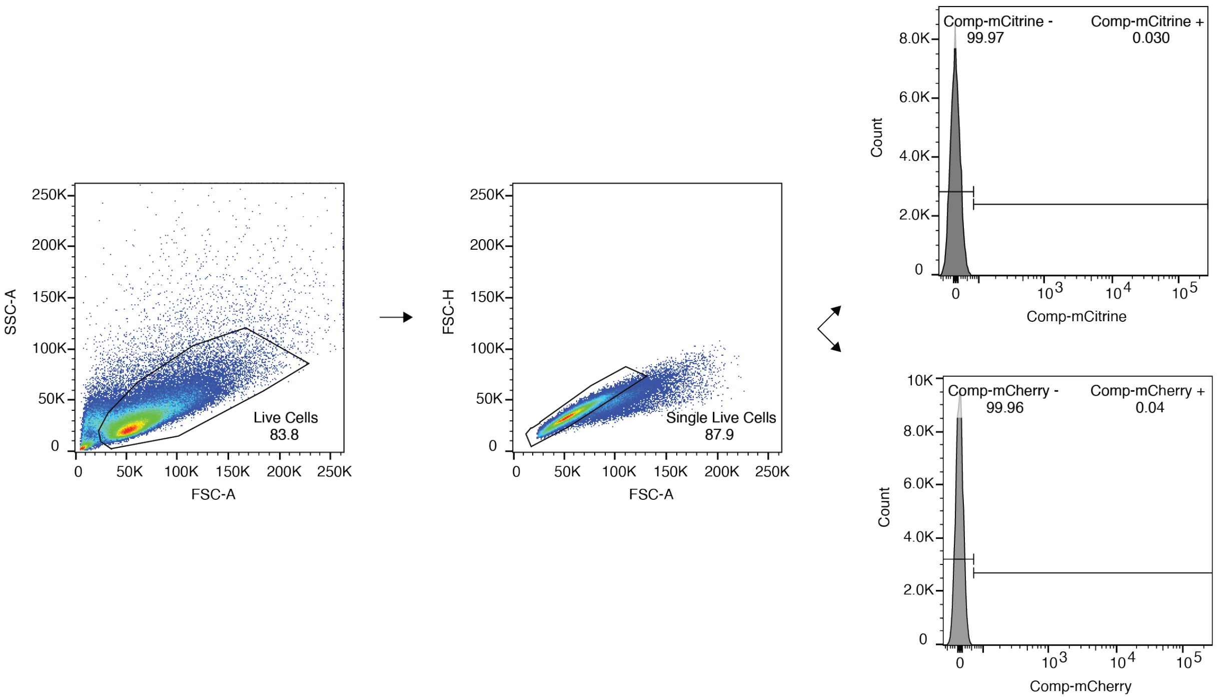


*Fig. S12: Cell gating strategy of HEK293 cells transfected with junk DNA*

Dot plots showing the strategy for gating: live cells (left) based on FSC-A vs SSC-A, live-single cells (middle) based on FSC-A vs FSC-H. Histograms of live-single cells in non-transfected cells (compensated) were used for gating positive and negative populations for the fluorescent channel of mCitrine (right,top) or mCherry (right,bottom).

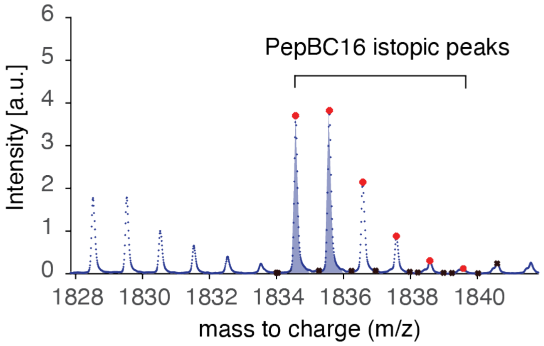


*Fig. S13: Calculation of peptide ion abundance from an MS spectrum*

A spectrum depicting the output of the automated MS spectrum analysis. The “peptide_auc_analysis” function identifies all peaks in a small m/z spectrum region corresponding to the m/z of PepBC16. Isotopic peaks assigned to PepBC16 are marked with a red dot, while impurity peaks are marked with a black sign. Any peaks outside of the PepBC16-associated spectrum segment are ignored. The AUC of the first two PepBC16 isotopic peaks are summed to calculate the PeBC16 ion abundance.
